# Disrupting the transmembrane domain interface between PMP22 and MPZ causes peripheral neuropathy

**DOI:** 10.1101/2023.12.24.573255

**Authors:** Natalya Pashkova, Tabitha A. Peterson, Christopher P. Ptak, Stanley C. Winistorfer, Debbie Guerrero-Given, Naomi Kamasawa, Christopher A. Ahern, Michael E. Shy, Robert C. Piper

## Abstract

PMP22 and MPZ are abundant myelin membrane proteins in Schwann cells. The MPZ adhesion protein holds myelin wraps together across the intraperiod line. PMP22 is a tetraspan protein belonging to the Claudin superfamily. Loss of either MPZ or PMP22 causes severe demyelinating Charcot-Marie-Tooth (CMT) peripheral neuropathy, and duplication of PMP22 causes the most common form of CMT, CMT1A. Yet, the molecular functions provided by PMP22 and how its alteration causes CMT are unknown. Here we find MPZ and PMP22 form a specific complex through interfaces within their transmembrane domains. We also find that the PMP22 A67T patient variant that causes a loss-of-function (Hereditary Neuropathy with Pressure Palsies) phenotype maps to this interface, and blocks MPZ association without affecting localization to the plasma membrane or interactions with other proteins. These data define the molecular basis for the MPZ~PMP22 interaction and indicate this complex fulfills an important function in myelinating cells.

## Introduction

Large diameter axons in the peripheral nervous system (PNS) are insulated by wraps of myelin sheath produced by Schwann cells to ensure rapid and efficient transmission of electrical impulses. Heritable neuropathies of the PNS are often referred to as Charcot Marie Tooth (CMT) disease. CMT is the most frequently diagnosed set of genetic neuro-muscular diseases, affecting 1:2500 individuals ^1^. Autosomal dominant (AD) inheritance CMT is the most common, followed by X-linked and autosomal recessive (AR) forms. Most forms of CMT are demyelinating and characterized by poorly formed myelin (CMT1), whereas one-third of CMT cases are primary axonal disorders (CMT2) in which axons degenerate but myelin formation is relatively normal ^2,3^.

Two crucial Schwann cell proteins involved in the structure and function of myelin are Myelin Protein Zero (MPZ/P0) and Peripheral Myelin Protein 22 (PMP22). Alteration of these proteins can cause CMT. MPZ comprises ~50% of myelin protein ^4,5^, and is a single-pass transmembrane protein with a single immunoglobulin-like (Ig) extracellular domain and cytosolic tail ^6,7^. It is post-translationally modified by N-linked glycosylation, sulfation, palmitoylation, and phosphorylation ^8,9^. MPZ is an adhesion protein, holding together adjacent wraps of myelin membrane across the intraperiod line ^8,10-13^. MPZ also uses its cytosolic tail to alter the lipid bilayer, and promote myelin compaction of the cytosolic region between myelin wraps to form the major dense line ^4,11,14-17^. Thus, MPZ resembles other adhesion molecules such as cadherins and integrins that couple their extracellular domains to the cytoskeleton as well as providing signaling cascades through their cytosolic domains. Variants in MPZ account for 5% of CMT cases overall and can cause demyelinating or axonal phenotypes, reflecting the diverse roles of MPZ in Schwann cells ^2,18,19^. Exactly how MPZ can bridge myelin wraps is not clear. For instance, the structural basis for how the extracellular Ig domain can adhere to an apposing MPZ Ig domain within compact myelin has not been fully resolved. A central concept for how MPZ works to form myelin is that it may oligomerize to form small foci within the membrane to increase its ability to adhere across the intraperiod line as well as alter the lipid bilayer to foster condensation of the cytoplasmic spaces within the major dense line. Evidence from crystallography studies and solution binding studies show that the extracellular domain of MPZ can oligomerize such that it would bring 4 MPZ proteins together in the plane of a membrane ^20-23^. Abundant membrane particles that correspond to the size of MPZ tetramers are evident in freeze-fracture electron microscopy studies on peripheral myelin ^24-26^. In addition, MPZ might be concentrated in small networks within lateral membrane ‘raft’ microdomains ^26 27-31^.

PMP22 is also a major component of myelin. There have been many functions proposed for PMP22, yet the molecular and cellular mechanisms that PMP22 uses to promote normal myelination remain unclear ^32-40^. PMP22 is a glycosylated 4-pass integral membrane protein that belongs to a subfamily of claudin-related proteins that includes EMP1, EMP2, and EMP3 ^41^. An emerging view is that the PMP22/EMP family of proteins promote adhesion and interact with adhesion proteins ^42-47^. Unlike MPZ, which is exclusively expressed in Schwann cells, PMP22 shows a broader pattern of expression and has been found at tight junctions between epithelial cells suggesting it can help organize adhesion structures ^4,40,48^. Moreover, PMP22 can help adhere membranes together in a variety of *in vitro* contexts ^49-51^. Consistent with a potential role in adhesion, PMP22 has been found to associate (directly or indirectly) with a variety of other adhesion proteins including Jam-C, integrin, myelin-associated glycoprotein (MAG) and oligomerization with PMP22 itself ^52-55^. Overall, alteration of PMP22 accounts for approximately 50% of CMT cases ^56,57^. The majority of demyelinating cases of CMT (CMT1A) are caused by an extra copy of wildtype PMP22 ^58-61^. However, heterologous deletion of PMP22 causes HNPP (Hereditary Neuropathy with Pressure Palsies), a milder form of peripheral neuropathy reflecting a partial loss-of-function phenotype ^62^. These observations underscore the importance of having the critical balance of PMP22 levels for proper myelination. Most newly synthesized PMP22 is degraded in Schwann cells ^63^, and part of how overexpression or variants of PMP22 cause CMT may be due to Endoplasmic Reticulum (ER) stress ^59,61,64-66^, exacerbated by PMP22’s ability to alter ER membrane morphology ^49^. However, whether that serves as the primary mechanism for CMT pathogenesis or whether PMP22 works in conjunction with other proteins to cause CMT is not clear.

We found that MPZ forms a strong and specific complex with PMP22. Though previous studies suggested MPZ forms a complex with PMP22 ^50,67^, the biochemical basis and specificity of that interaction were not defined nor validated by others. Our data demonstrate a robust and specific interaction mediated by their transmembrane domains (TMD). Within these interacting interfaces lies the patient variant A67T, which causes an HNPP phenotype. We find that the A67T variant disrupts formation of the MPZ~PMP22 complex without affecting PMP22 localization to the plasma membrane nor association with other PMP22-interacting proteins. These data suggest the interaction between MPZ and PMP22 serves a critical function in myelin.

## Results

### MPZ forms a complex with PMP22

We found that PMP22 forms a very specific and strong interaction with MPZ in HEK293 cells and rat RT4 Schwannoma cells. Figure 1 shows that PMP22-GFP co-immunoprecipitates C-terminally HA-tagged MPZ when both are co-expressed in HEK293 cells (Figure 1A) and RT4 Schwannoma cells (Figure 1C). However, no complex was found between MPZ and the PMP22-related protein EMP3 (Figure 1C). Immunoprecipitations were accomplished using Sepharose coupled to α-GFP nanobodies. In general, these observations partly align with previous studies showing evidence that PMP22 and MPZ form a complex ^50,67^. We found non-ionic detergents destroyed the complex whereas a detergent formulation of DDM (n-dodecyl-β-d-maltoside) and CHS (cholesterol-like cholesteryl hemisuccinate) preserved the complex (Figure 1A). Previous studies used a mix of non-ionic detergents within their lysis buffer (NP40 and Triton X-100)^67^. In contrast, other detergents that better preserve structure and function have been identified; these include DDM, which is a mild non-ionic detergent that is typically efficient at solubilizing membrane proteins ^68,69^. Often, the best preservation of membrane protein structure is achieved with the additional inclusion of CHS ^70,71^. Previous studies indicate that PMP22 is cholesterol-philic and that higher-order oligomers of PMP22 *in vitro* appear to be preserved using cholesterol-like environments ^29,38,39,54,72,73^. These biophysical properties provide a foundation to study the MPZ~PMP22 complex and explain why detecting this complex may have been difficult for other investigators. We also found that DDM/CHS was as effective in solubilizing available MPZ and PMP22 from HEK293 cells as 1% SDS, showing that DDM/CHS solubilizes each protein efficiently and does not work on a subpopulation (Figure S1A). In addition, the inclusion of CHS to DDM greatly enhanced the ability to co-immunoprecipitate MPZ-HA with PMP22-GFP indicating that cholesterol may help drive their association (Figure S1B).

**Figure 1:**
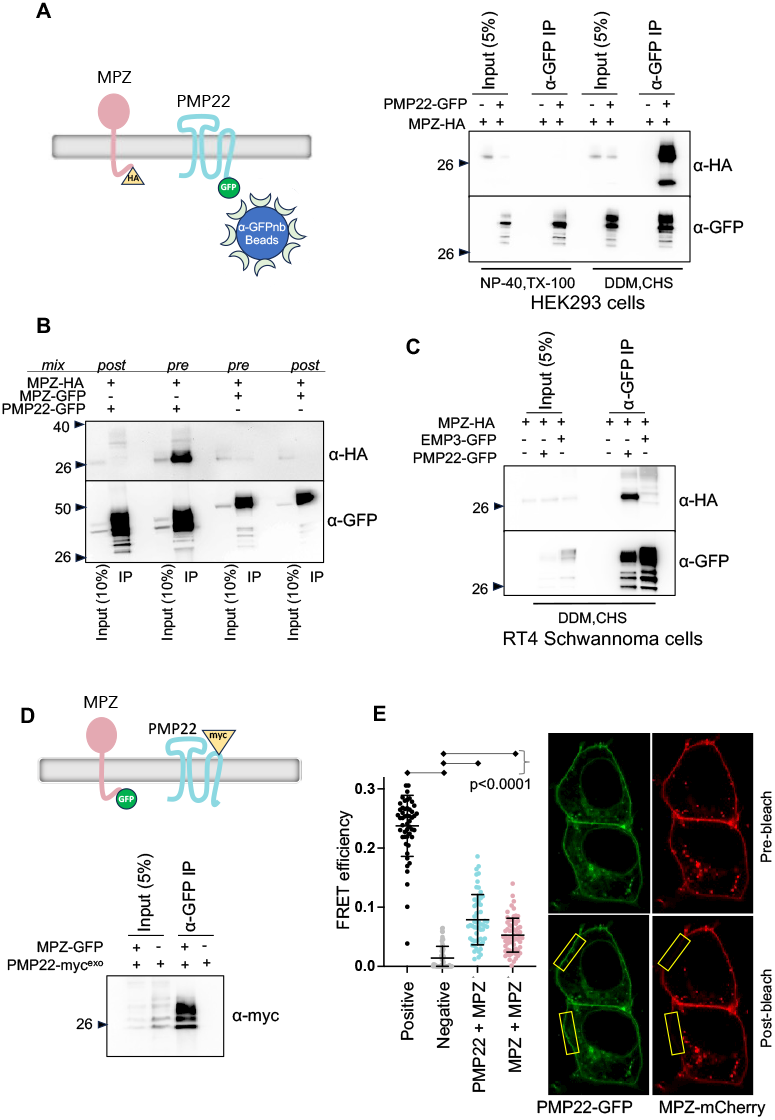
MPZ and PMP22 form a complex in HEK293 and RT4 Schwannoma cells. **A**. Left panel shows scheme for co-immunoprecipitation in which MPZ carrying a C-terminal HA eiptope tag and PMP22 carrying a C-terminal GFP tag are co-expressed and then immunoprecipitated with Sepharose linked to α-GFP nanobodies. Right panel shows results of co-immunoprecipitation. Lysates from HEK293 cells transiently co-transfected with MPZ-HA and PMP22-GFP were prepared with 0.5% NP-40 and 0.5% Triton X-100 or with 1% DDM and 0.2% CHS detergents. Bead-bound precipitates were washed in PBS with their respective detergents and eluted with SDS sample buffer. A proportion of the input (5%) was analyzed along with precipitated complexes by SDS-PAGE and immunoblotting with mouse α-HA (to detect associated MPZ-HA) and mouse α-GFP (to show efficiency of PMP22-GFP immunoprecipitation). **B.** The co-immunoprecipitation scheme in A was used for HEK293 cells in which PMP22-GFP and MPZ-HA were co-expressed together before (pre) lysis or were expressed in separate populations and mixed after lysis (post). Co-immunopreciptiation was measured between PMP22-GFP and MPZ-HA or MPZ-GFP and MPZ-HA. Immunoprecipitates together with a 10% equivalent of the starting input was immunoblotted with anti-HA to reveal associated MPZ-HA. Note that while MPZ-HA associates with MPZ-GFP, the level of this association is far less than that with PMP22-GFP. Moreover, both associations required that proteins had to be expressed in the same cells prior to detergent lysis to detect the presence of MPZ~MPZ or MPZ~PMP22 complexes. **C**.Co-immunoprecipitation of PMP22-GFP and MPZ-HA from DDM/CHS detergent lysates of transiently transfected RT4 Schwannoma cells. Whereas MPZ-HA was co-precipitated with PMP22-GFP, it was not co-precipitated with the PMP22-related protein EMP3-GFP. **D**. Top panel shows scheme for co-immunoprecipitation of MPZ-GFP with PMP22 containing an exofacial myc epitope tag inserted into the second extracellular loop. Bottom panel shows PMP22-myc^exo^ is recovered when co-expressed with MPZ-GFP and immunoprecipitated with α-GFP nanobody beads. **E**. Quantitation of FRET efficiency between PMP22-GFP and MPZ-mCherry and between MPZ-GFP and MPZ-mCherry. FRET was measured as an increase in donor fluorescence after acceptor photobleaching in transiently transfected HEK293 cells. The ‘positive’ control was MPZ tandemly fused to both msGFP2 and mCherry. The ‘negative’ control was Glycophorin A-GFP and MPZ-mCherry. Regions of interest were analyzed and quantified for at least 25 cells from 2 different experiments. FRET efficiencies were calculated with the Leica LAS X Microlab module. All ROI data are represented singly and aggregated as mean FRET efficiency ± S.D (from left to right 0.237±0.052, 0.014±0.020, 0.079±0.042, 0.053±0.029). One-way Anova with Turkey’s multiple comparison test revealed the negative control was significantly different to each of the other samples. On the right is a representative FRET image of PMP22-GFP + MPZ-mCherry. The yellow boxes on the images show regions of interest used for bleaching and FRET measurements.

To investigate these interactions, we used GFP-tagged PMP22 where GFP was fused to the PMP22 C-terminus. Previous studies found that GFP fusion to PMP22 had the potential to drive aggregation ^74-76^, which could confound our co-immunoprecipitation experiments. We sought to circumvent this problem by using monomeric superfolder GFP (msGFP2) ^77^ separated by a short flexible linker. As tested by size exclusion chromatography, DDM/CHS-solubilized PMP22-GFP eluted as a major peak coincident with an apparent molecular weight of ~150-300kDa compared to size exclusion chromatography standards. No significant proportion of heterogeneous GFP-fusion aggregates were found in the void volume resolved by Superose 6 column indicating PMP22-GFP did not aggregate (Figure S1C). To confirm the MPZ~PMP22 interaction, we also used a PMP22 carrying a myc tag inserted into its second extracellular loop, previously reported to provide an epitope-tagged form of PMP22 that is not prone to aggregation ^74^. Figure 1D shows that PMP22-myc^exo^ also efficiently co-immunoprecipitated with MPZ-GFP. We also examined the MPZ~PMP22 complex by size exclusion chromatography of DDM/CHS lysates from HEK293 cells co-expressing PMP22-GFP and MPZ-HA. When expressed together, each protein was found absent from large aggregates in the void volume. Moreover, the complex, revealed by immunoprecipitating PMP22-GFP across several eluted fractions, was also not found as a large aggregate and migrated with an apparent MW of ~200-350 kDa (Figure S1D).

We next determined whether the PMP22~MPZ complex can only be formed from proteins expressed together in the same cells. Figure 1B shows that when MPZ-HA and PMP22-GFP are co-expressed in the same cell (labeled as ‘pre’), they can efficiently be co-immunoprecipitated. However, when expressed in separate cell populations and mixed after lysis (labeled as ‘post’), the PMP22~MPZ complex could not be recovered by co-immunoprecipitation. In all experiments, we found that a large proportion of MPZ-HA was in PMP22-GFP immunoprecipitates, indicating that the association of MPZ with PMP22 is strong, efficient, and may be near stoichiometric. We also find that MPZ-HA forms a complex with MPZ-GFP when expressed in the same cell, consistent with its ability to oligomerize ^20,78,79^. In contrast, however, the efficiency of MPZ~MPZ complex recovery was far less than that of the PMP22~MPZ complex recovery (Figure 1B).

In addition, we confirmed that PMP22-GFP associated with MPZ-mCherry within the plasma membrane of HEK293 cells as assessed by FRET measured by donor fluorescence-enhancement after acceptor photobleaching (Figure 1E). MPZ tandemly fused to both msGFP2 and mCherry was used as a positive control which gave a percent FRET efficiency of 23.7%. As a negative control we used MPZ-mCherry co-expressed with glycophorin A-GFP, both of which localize to the plasma membrane but are not known to interact. The combination of MPZ-mCherry with PMP22-GFP had a FRET efficiency significantly different than the negative control pair (7.9%±4.2 vs 1.4%±2.0, respectively). Co-expressed MPZ-mCherry and MPZ-GFP also had a significant FRET efficiency above the negative control (5.3%±2.9 vs 1.4%±2.0, respectively). Most ROIs used for FRET measurements were areas of the plasma membrane that were not juxtaposed to other cell plasma membranes. For the few ROIs that we sampled on the plasma membrane areas between 2 cells, we saw no increased FRET efficiency. All of which indicated that the FRET signal we observed was from proteins interacting within the same bilayer in *cis*.

### Specificity of the MPZ~PMP22 association

Both MPZ and PMP22 each belong to subfamilies of homologous proteins (Figure S2). We found MPZ is the only member of its subfamily to interact with PMP22, and PMP22 is the only member of its subfamily to interact with MPZ. MPZ belongs to a subfamily of single-pass transmembrane proteins, with extracellular Ig-like domain containing adhesion proteins including MPZL1, MPZL2, and subunits of voltage-gated sodium channels (SCN2B, SCN3B,and SCN4B) ^80^. Figure 2A shows that PMP22 strongly co-immunoprecipitated with MPZ-HA, very weakly with MPZL2, and showed no co-immunoprecipitation with other MPZ family members MPZL1, SCN2B, SCN3B, or SCN4B. In contrast, while the PMP22 related protein EMP3 did not interact with MPZ-HA, it did interact strongly with MPZL2, SCN2B and SCN3B and also had a weak interaction with MPZL1.

**Figure 2.**
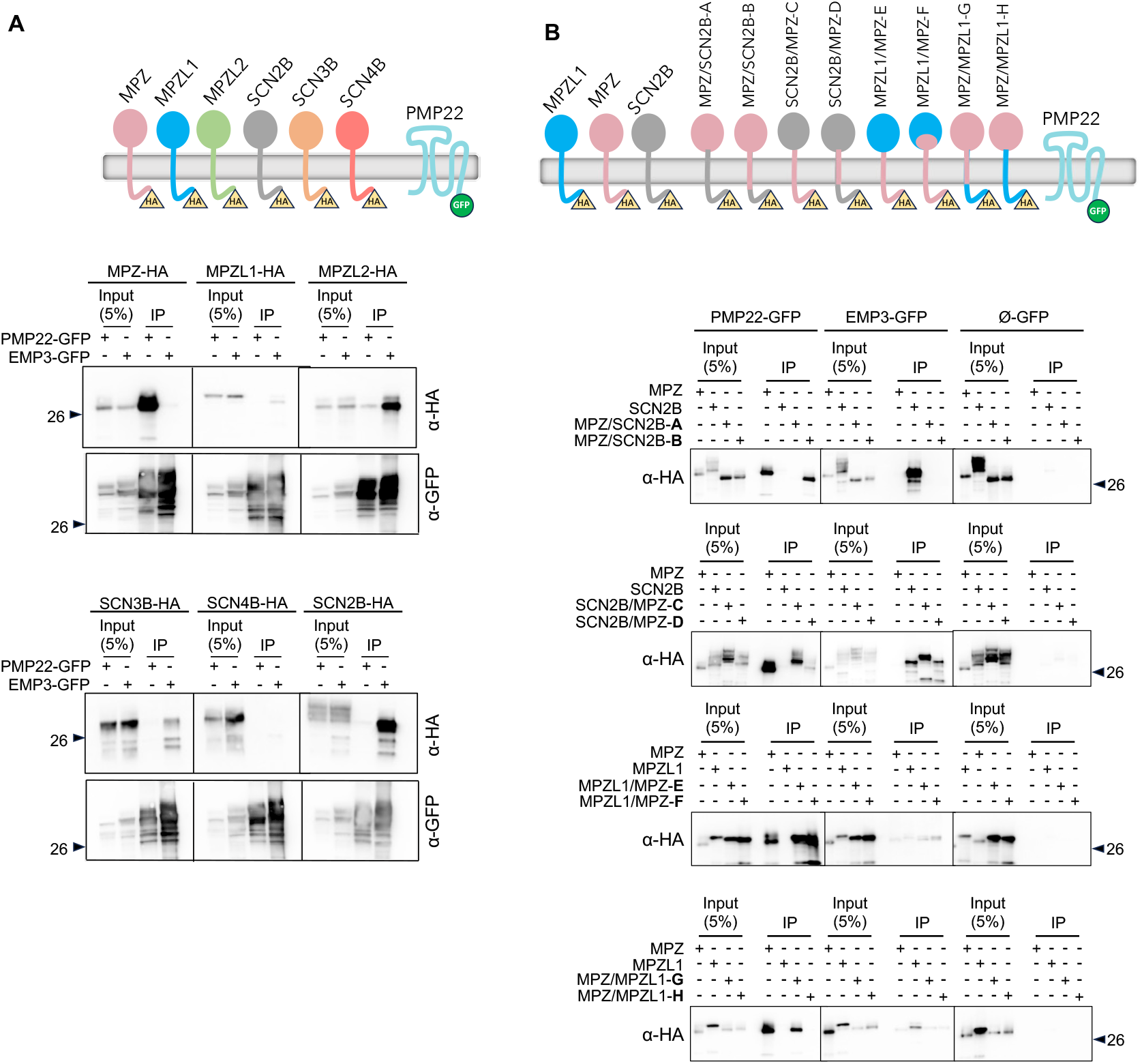
Specificity of MPZ association with PMP22. **A**. Several members of the MPZ/β-subunit family were assessed for their ability to associate with PMP22-GFP using the co-immunoprecipitation scheme outlined in Figure 1. DDM/CHS detergent lysates from HEK293 cells expressing the indicated HA-tagged MPZ/β-subunit proteins with either PMP22-GFP or EMP3-GFP were immunoprecipitated with α-GFP nanobody beads. Immunoprecipitation of PMP22-GFP only efficiently recovered MPZ-HA and a very low level of MPZL1. In contrast, MPZL2, SCN3B, and SCN2B formed strong complexes with EMP3-GFP. A 5% equivalent of the input lysate was included in each immunoblot. **B**. The indicated HA-epitope tagged chimeric proteins made of the extracellular Ig-like domain, TMD, and cytosolic domains of MPZL1, SCN2B, and MPZ were assessed for association with PMP22-GFP, EMP3-GFP, or GFP alone (Ø-GFP) as described in A. Chimeras containing the TMD of MPZ were all co-precipitated with PMP22-GFP whereas substitution of the extracellular domain or cytosolic domain of MPZ with that of SCN2B or MPZL1 did not block PMP22 association. In contrast, the ability of SCN2B to associate with EMP3-GFP is conferred by its extracellular Ig-like containing domain.

We next exploited this specificity to map the regions necessary and sufficient for PMP22~MPZ interaction using a set of chimeric MPZ/β-subunit proteins where the extracellular Ig-like domain, transmembrane domains, and cytosolic tails were swapped. The exact composition of these chimeric proteins is provided in supplemental data, described in the key resources table, and provided schematically in Figure 2B. Chimeras made of parts of MPZL1 and SCN2B that contained the MPZ TMD were all able to form a complex with PMP22-GFP as assessed by co-immunoprecipitation. These include chimeras B, C, and D (containing the MPZ TMD within the context of SCN2B) and chimeras E, F and G (containing the MPZ TMD within the context of MPZL1). We also examined whether these chimeras were informative about the regions that mediate association of EMP3 with SCN2B. Here, the data indicated that the extracellular Ig-like domain-containing region of SCB2B was necessary and sufficient for interaction with EMP3. Together, these data emphasize the specificity that PMP22 has for associating with MPZ over other MPZ/β-subunit family members and demonstrate that association is mediated through the MPZ TMD. These data also show that association of other MPZ/β-subunit members with different PMP22 family members (e.g. SCN2B association with EMP3) is not exclusively through transmembrane regions.

### PMP22 Interaction surface on the transmembrane domain of MPZ

We next determined which amino acids within the MPZ TMD were critical for PMP22 association. Because TMD interactions are mediated primarily via shape complementarity ^81^, we altered large bulky hydrophobic residues to small ones (e.g. I/L to A) and *vice versa*. These mutations were made in the context of MPZ-mCherry to allow the subcellular localization of each mutant to be easily followed by microscopy to ensure that loss of PMP22 interaction was not due to severe misfolding, as would be indicated by retention in the ER. Each MPZ-mCherry mutant was co-expressed with PMP22-GFP and tested for association by co-immunoprecipitation with α-GFP nanobody beads. Figure 3A shows that mutations in a subset of MPZ TMD residues disrupted association with PMP22. These residues largely clustered on one face of the predicted transmembrane α-helix and located on the half of the TMD closest to the extracellular side of the membrane (Figure 3D). This cluster contains residues G155, L158, and L162, that we further confirmed as critical for interaction in the context of a C-terminal HA epitope-tagged MPZ (Figure 3B). We also found that neither these mutations singly nor in combination had deleterious effects on the ability of MPZ-mCherry to traffic to the plasma membrane (Figure 3C) nor the ability to form an oligomeric complex with MPZ-GFP (Figure 3B and Figure S3A). Together, these data confirm that the interaction with PMP22 is conferred through the MPZ TMD and highlight a discrete region within this domain that is likely to comprise the binding interface. In addition, these data provide well-characterized mutant versions of MPZ for *in vitro* and *in vivo* functional studies that have specifically lost their ability to associate with PMP22 but retain other protein interactions as well as correct trafficking to the cell surface.

**Figure 3.**
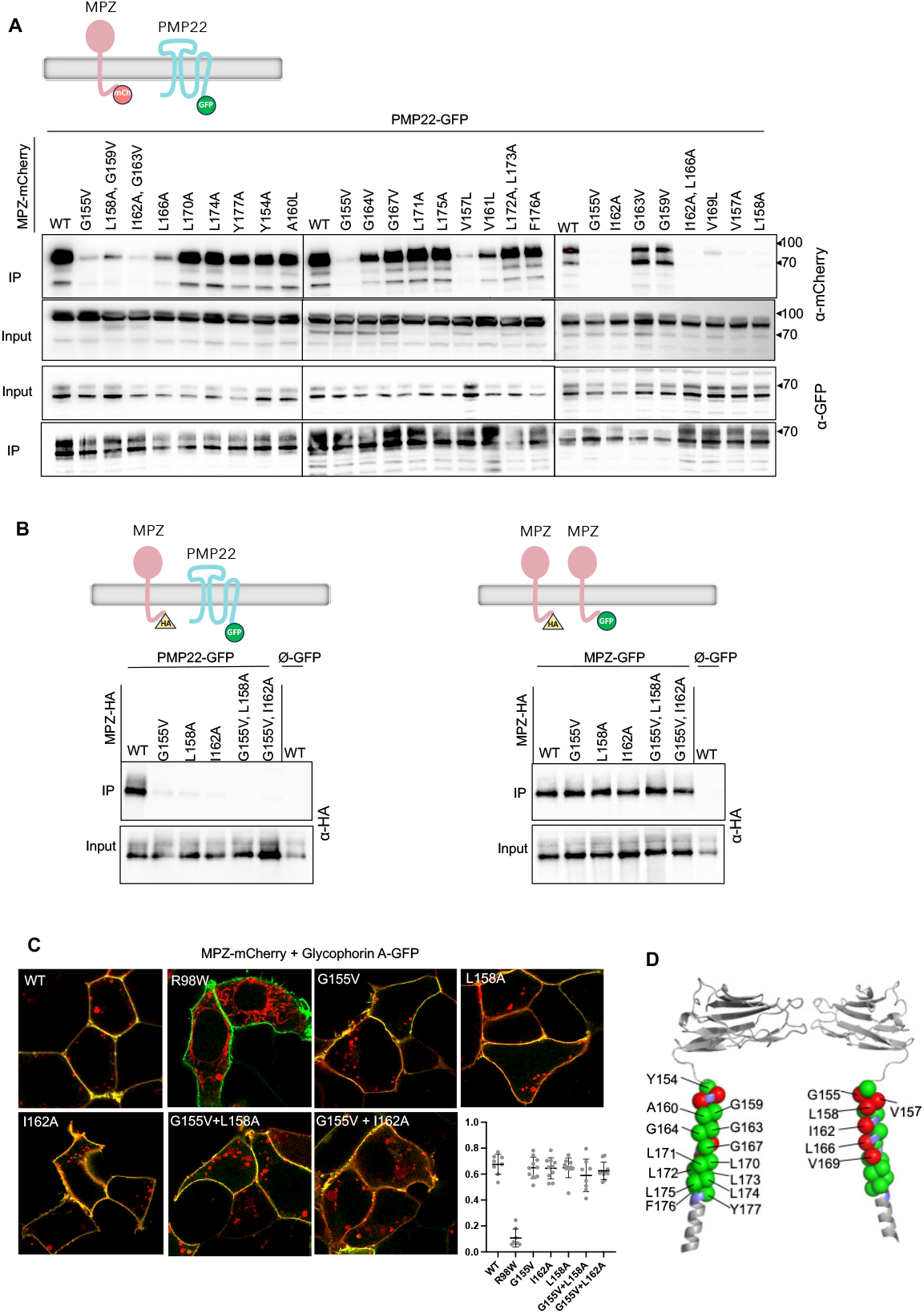
Mapping the PMP22-interacting interface within the MPZ transmembrane segment. **A**. Top panel shows experimental scheme in which PMP22-GFP is co-expressed with mutant versions of MPZ-mCherry and immunoprecipitated with α-GFP nanobody beads. Lower panel shows immunoprecipitation results on the indicated MPZ-mCherry mutants. **B**. Left. Results from A were verified using MPZ mutants in the context of a MPZ C-terminally HA-epitope-tagged protein co-expressed with PMP22-GFP or GFP alone (Ø-GFP) by coimmunoprecipitation with α-GFP nanobody beads and immunoblotting with α-HA antibodies. Right. WT and the indicated MPZ-HA mutants were assessed for their ability to oligomerize with MPZ-GFP using α-GFP nanobody beads. **C**. Cell surface localization of WT MPZ-mCherry and the indicated mutants was assessed by confocal microscopy. Micrographs show WT MPZ-mCherry at the plasma membrane marked by Glycophorin A tagged with GFP, and puncta likely corresponding to endosomes. The MPZ R98W mutant, which is known to be retained in the ER, was included for comparison. Multiple fields encompassing >100 cells were assessed for overlap of MPZ-mCherry with the plasma membrane marker Glycophorin A using Manders overlap coefficient and plotted for each field in graph at right along with mean +SD. One way ANOVA revealed no differences in cell surface localization (p>0.1) amongst mutants and the WT MPZ-mCherry with the exception of the R98W mutant that had a significantly lower cell surface localization (p<0.0001) to every other MPZ-mCherry protein analyzed. **D**. Residues in the MPZ TMD that when mutated had no effect (green) and dramatic reduction (red) in the ability to associate with PMP22 were mapped onto a model (AlphaFold: AF-P25189-F1) of a portion of MPZ containing the extracellular Ig-like domain and the α-helical region encompassing the TMD. Note that the red residues align along one face of the α-helix forming a cluster indicating the likely interface mediating PMP22 association.

### Basis for how PMP22 interacts with MPZ

Figure 4A shows that among the PMP22/EMP family of proteins, only PMP22 formed a complex with MPZ whereas GFP-tagged EMP1, EMP2 or EMP3 did not. To find regions of PMP22 that contribute to MPZ association, we initially made a set of chimeric proteins between PMP22 and EMP3 for co-immunoprecipitation experiments. However, this approach was uninformative as chimeras containing swaps of 1 to 3 different transmembrane helices or extracellular loops largely compromised the ability of these chimeric proteins to associate with MPZ and did not yield a clear pattern for what region was required for association. Based on the data in Figure 3 indicating that residues residing in one half of the MPZ TMD were important for PMP22 association, we next focused on amino acid substitutions within the PMP22 transmembrane domains concentrating on residues within the four membrane-spanning helices closest to the extracellular side. Rather than use patient variants, we systematically analyzed each transmembrane helix by altering large hydrophobic residues to small ones (and vice versa) and assessing the impact on co-immunoprecipitation of these mutant PMP22 proteins with MPZ (Figure 4B). To rationalize this mutagenesis strategy, we mapped these amino acid substitutions onto a previously determined homology model for PMP22 that was made based on its similarity to claudin-15 (PDB:4P79) for which a crystal structure is known ^82,83^. We also used a model of PMP22 generated by Alphafold ^84^ (Figure 4C), which predicted a structure similar to claudin-15 and the previously determined PMP22 homology model (RMSD 0.98 and 1.311, respectively). Importantly, both models were highly similar within the membrane spanning segments (RMSD 0.95), bolstering the prediction of which residues were likely facing out of the TMD helical bundle and positioned for potential interactions with another protein versus those residues with side chains making intramolecular interactions between transmembrane helices that would likely be critical for the integrity of the PMP22 structure. Accordingly, we focused our mutagenesis on residues predicted to face outwardly. Several amino acid substitutions in PMP22 were found to ablate or dramatically reduce interaction with MPZ (Figure 4B,C). These include the I70A, L71A, F75A amino acid substitutions that map to a cluster of residues on one face of the 2^nd^ transmembrane helix of PMP22 and blocked MPZ association altogether. Altering residues near this cluster (e.g. I73A and I74A) but on a different face of the second transmembrane helix diminished but did not destroy interaction with PMP22. Alteration of other residues along the 2^nd^ transmembrane helix of PMP22 more toward the cytosolic region did not affect MPZ association. Similarly, amino acid substitutions within the 1^st^, 3^rd^, and 4^th^ transmembrane helices of PMP22 were without effect on MPZ association with the exception of I116A near the extracellular edge on the 3^rd^ transmembrane helix. Finally, while wildtype MPZ and PMP22 interact as assessed by FRET within the plasma membrane of HEK cells, MPZ^G155V^ and PMP22^F75A^ did not produce a significant FRET signal when co-expressed, confirming that these mutations disrupt interaction within cells (Figure S3B).

**Figure 4.**
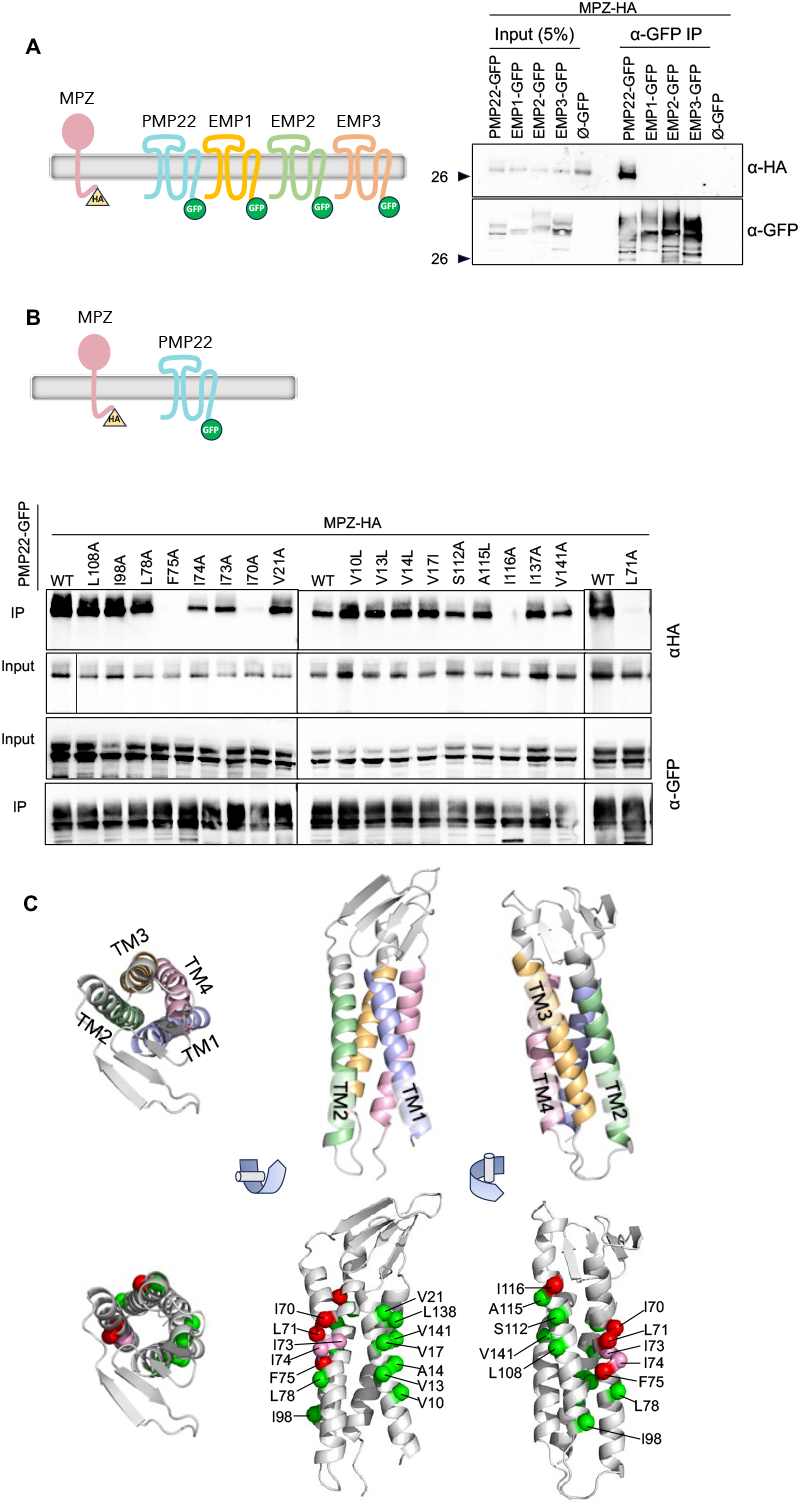
Specificity of PMP22 interaction with MPZ. **A**. PMP22-GFP or the PMP22-related proteins EMP1-GFP, EMP2-GFP, EMP3-GFP, or GFP alone (Ø-GFP) were co-expressed with MPZ-HA in HEK293 cells and immunoprecipitated with α-GFP nanobodies. Immunoprecipitated eluted proteins were immunoblotted with α-HA to detect associated MPZ-HA and α-GFP to monitor recovery of PMP22-GFP. A 5% equivalent of the input lysate was included in the immunoblots. **B**. The indicated amino acid substitutions were incorporated into PMP22-GFP and assessed for association with co-expressed MPZ-HA. **C**. Model of PMP22 (AlphaFold: AF-Q01453-F1) showing orientation of the four transmembrne helices (TM1-4) of PMP22 and the predicted position of the amino acids critical for MPZ binding. Residues in PMP22 that when mutated had no effect (green), a partial reduction (pink) and dramatic reduction (red) in the ability to associate with MPZ are shown.

The residues identified as disrupting the interaction between MPZ and PMP22 were used as restraints to generate a docking model that describes how MPZ could form a complex with PMP22 (Figure 5). Here, residues on one face of the 2^nd^ transmembrane helix of PMP22 (I70, L71, and F75) are docked to residues on one face of the MPZ TMD (G155, L158, I162, and L166). It is not apparent in this model for how alteration of residue I116 to Alanine on the 3^rd^ transmembrane helix of PMP22 blocks association with MPZ since I116 does not interact directly with MPZ. In Alphafold models of PMP22, I116 is packed against transmembrane helix 2 (< 3 Å to W61, V65; < 4 Å to S64, T68; < 5 Å to L62) and the I116A mutation leads to a 43% reduction in residue-residue van der Waals contacts between the exofacial halves of the 2^nd^ and 3^rd^ transmembrane helices as assessed on the RING webserver ^85^. The position of I116 suggests that a mutation may act indirectly by introducing structural or dynamic changes that alter the ability of the 2^nd^ PMP22 transmembrane helix to bind MPZ. Alternative models that include I116 as a docking restraint (shown in Data Figure S4) show a loss or reduction of a number of experimentally predicted residue interactions and a correspondingly higher restraints violation energy. These docking models are speculative and rely on the accuracy of the predicted structures for the individual interacting partners. Nonetheless, they are instructive for highlighting potential interacting surfaces within the TMDs of PMP22 and MPZ that our mutagenesis and co-immunoprecipitation experiments define.

**Figure 5.**
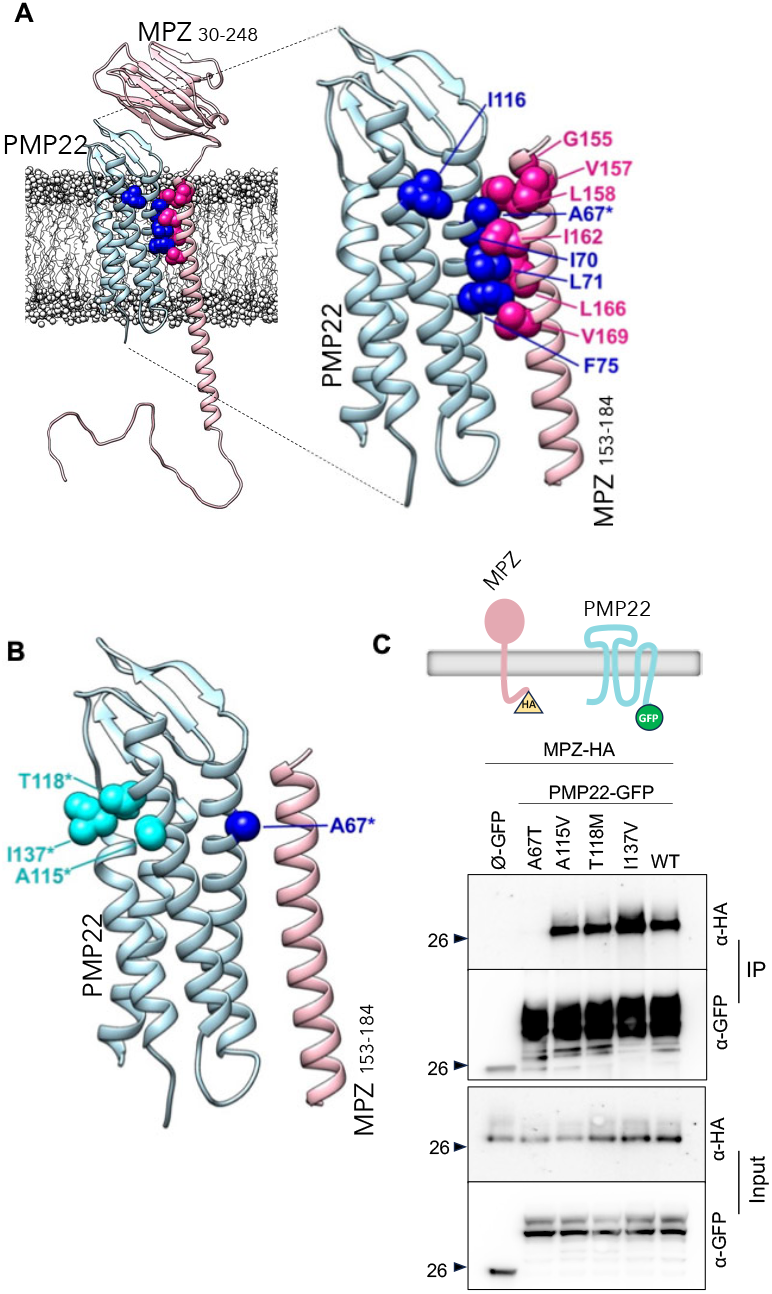
The patient variant of PMP22 (A67T) causing an HNPP phenotype maps to the presumed interface between PMP22 and MPZ. **A**. Model of the complex between PMP22 and MPZ in which association is mediated by residues identified in amino acid substitution experiments. (Upper panel) The multi-component model composed of the full-length human PMP22 (light blue – AlphaFold: AF-Q01453-F1) and MPZ (pink – AlphaFold: AF-P25189-F1) is oriented in a lipid bilayer of 60% POPC : 40% cholesterol (white). For a few unstructured residues between MPZ domains, torsion adjustments were made for improved visualization of the extracellular domain and the flexible portion of the cytosolic tail. The residues that destroy or diminish the association of PMP22 and MPZ when mutated are indicated. The position of the HNPP-causing patient variant, A67T, which lies within the predicted interface mediating MPZ~PMP22 binding is indicated (*). **B**. Model of the PMP22~MPZ interface shown in A with the positions of select patient variants including the A67T indicated on the different transmembranehelices **C**. Effect of PMP22 patient variants on the association with MPZ. MPZ-HA was co-expressed with GFP alone (Ø-GFP), WT PMP22-GFP or the indicated PMP22 mutants and assessed for complex assembly by coimmunoprecipitation with α-GFP nanobody beads and immunoblotting for MPZ-HA with α-HA antibodies. Comparable expression levels of the different PMP22 mutants as well as the efficiency was assessed by immunoblotting input lysates and bead immunoprecipitates with α-GFP antibodies.

### The interaction of PMP22 with MPZ is important for PMP22 function

We next surveyed whether known PMP22 amino acid substitution variants that cause a range of peripheral neuropathy phenotypes had an impact on complex formation with MPZ. We focused on residues predicted to be on the outer-facing surfaces of the TMDs and excluded residues that would potentially mediate intramolecular interactions between transmembrane helices and thus may be critical for proper PMP22 folding ^82^. The A67T patient variant is located in the 2^nd^ transmembrane helix in the midst of the cluster of residues we found by mutagenesis to be important for MPZ association (Figure 5A, B). An in silico PMP22 A67T substitution in our PMP22~MPZ docking model leads to atomic clashes with both L158 and I162 of MPZ suggesting that sidechain rearrangements and helix repositioning would be required to maintain intermolecular interactions. Further, A67T is predicted by SSIPe ^86^ to decrease the ΔΔG of binding by 0.343 kcal/mol, and by HADDOCK to decrease the buried interaction surface area when PMP22 A67T is docked to the TM helix of MPZ. Figure 5C shows that this patient variant dramatically blocked association with MPZ-HA as assessed by co-immunoprecipitation. In contrast, other PMP22 patient variants that are predicted to map to outwardly oriented residues had no effect on MPZ association. These included A115V and T118M in the 3^rd^ transmembrane helix and I137V in the 4^th^ transmembrane helix of PMP22. The A67T variant was described in a patient with an HNPP phenotype, reflecting a heterozygous loss of PMP22 function ^87^. This indicates that the association of PMP22 with MPZ may be a critical requirement for PMP22 to fully function *in vivo*.

Previous experiments indicate that several biophysical properties as well as trafficking efficiency to the cell surface is largely preserved in the PMP22 A67T patient variant compared to wild-type (WT) PMP22 ^73,88,89^. These observations support the idea that the loss of function for the A67T variant is specifically due to loss of MPZ association rather than an indirect effect due to perturbation of other properties. To test this idea further, we assessed whether PMP22 A67T preserved its ability to interact with other proteins (Figure 6). PMP22 has been found to form a complex with itself and with the double Ig-domain containing adhesion protein Jam-C ^52,55,90^. In addition, during the course of our experiments, we discovered that PMP22 also interacts with the Jam-B adhesion protein and the PMP22-related protein, EMP1. Therefore, we determined whether the A67T patient variant as well as other amino acid substitutions within the presumed interface of PMP22 that mediates MPZ association preserved these protein interactions. Figure 6A shows that the PMP22-GFP carrying the A67T substitution or the I70A, I74A, or F75A were able to oligomerize with PMP22-HA and EMP1-HA as determined by co-immunoprecipitation. Likewise, these PMP22 mutants were also able to co-immunoprecipitate both Jam-C-HA or Jam-B-HA (Figure 6B), demonstrating that other protein interactions are unperturbed by these amino acid substitutions and indicate the overall integrity of the PMP22 structure remained intact.

**Figure 6.**
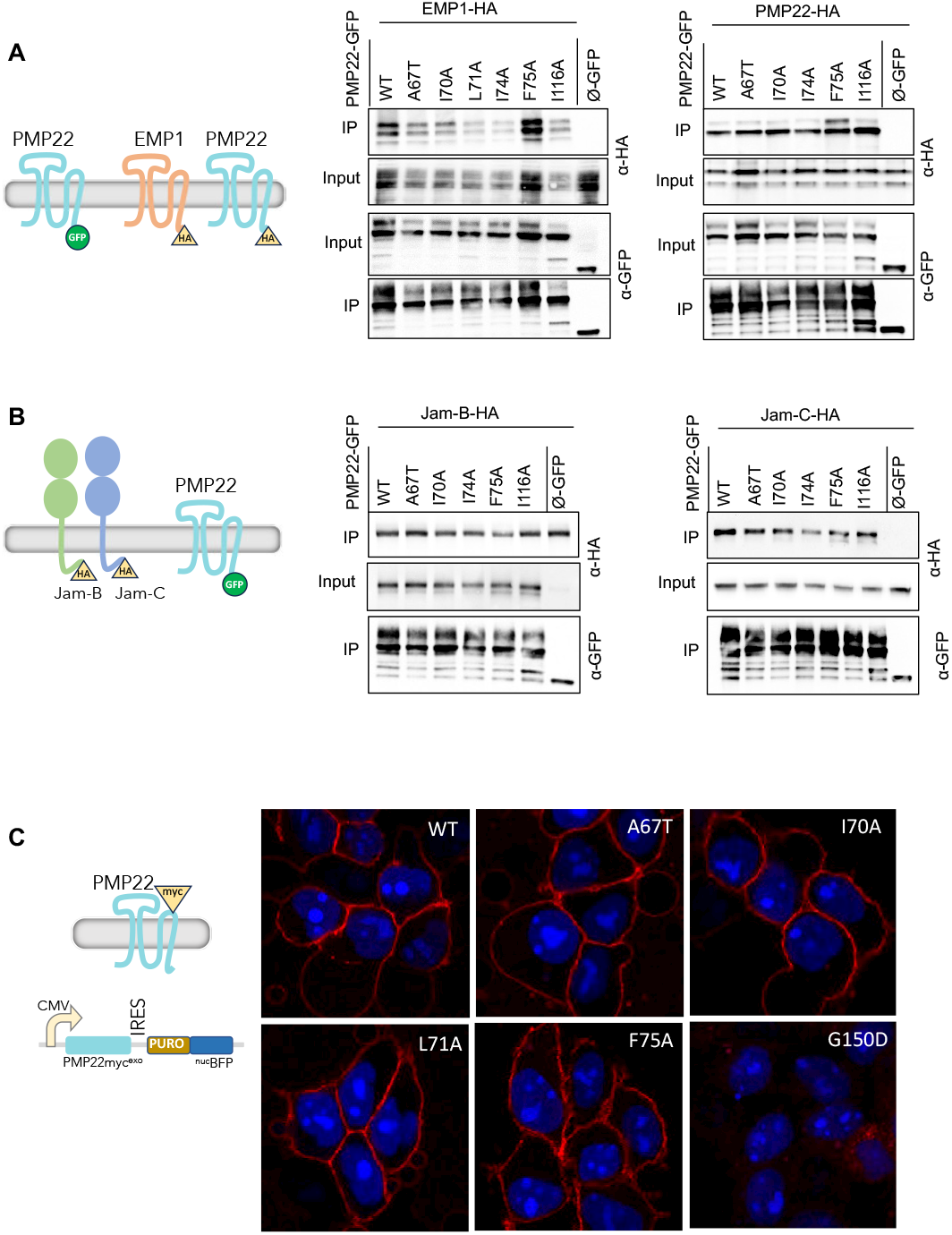
Specificity of PMP22 mutants lacking the ability to bind MPZ. **A**. PMP22-GFP, both WT and the indicated mutants along with GFP alone (Ø-GFP), were assessed for their ability to bind EMP1-HA (left) or PMP22-HA (right). Comparable expression levels of the different PMP22 mutants as well as the efficiency was assessed by immunoblotting input lysates and bead immunoprecipitates with α-GFP antibodies. **B**. PMP22-GFP, both WT and the indicated mutants, were assessed for the ability to bind Jam-B-HA (left) and Jam-C (right). **C**. The cell surface localization of WT and the indicated mutants of PMP22 was assessed in the context of the PMP22-myc^exo^ protein containing an extracellular (exofacial) myc epitope. Expression of PMP22-myc^exo^ was coupled to the expression of a nuclear-localized puromycin-blue fluorescent protein fusion by having it downstream of an internal ribosome entry site on the PMP22 mRNA. Stable transfected HEK293 cell lines expressing these mutants were fixed, left unpermeablized and labelled with α-myc monoclonal antibody and Alexa 568 secondary antibody prior to visualization by confocal microscopy (right).

We also measured how well each PMP22 mutant trafficked to the cell surface using a modified assay previously described that follows the exposure of a myc epitope tag when PMP22-myc^exo^ is localized to the cell surface ^65^. Here, PMP22-myc^exo^ is co-expressed on the same mRNA with a modified blue-fluorescent protein localized to the nucleus, providing a fiduciary signal to monitor the levels of PMP22-myc^exo^ production (Figure 6C). As a comparison, we also measured the trafficking efficiency of PMP22 G150D, the variant found in Trembler(Tr) mice known to trap PMP22 within the ER ^91^. As a qualitative comparison, we examined cell surface labeling of the exofacial myc epitope *via* immunofluorescence (Figure 6C). We also performed a quantitative analysis via flow cytometry (Figure S5). We found that PMP22-myc^exo^ carrying the A67T patient variant causing an HNPP phenotype was trafficked to the cell surface to the same extent as wildtype PMP22. Moreover, the amino acid substitutions within the 2^nd^ transmembrane helix we found *via* systematic mutagenesis (e.g. I70A, L71A, and F75A) also retained their ability to traffic to the cell surface. In contrast, PMP22 with the Trembler G150D mutation had a much lower efficiency of delivery to the cell surface and was found abundantly in the ER when cells were permeabilized before immunolabeling (Figure S6).

### Distribution of PMP22 and MPZ within the plasma membrane

There is evidence in myelin that the adhesion protein MPZ may be concentrated within the membrane, working within a large particle that may contain 4 MPZ proteins and perhaps other myelin proteins ^24-26^, as well as having a favored distribution in membrane microdomains on the order of 200-1000 nm ^27^. Such microdomains might perhaps be driven by the assembly of cholesterol and sphingolipid rich areas that form the biochemical basis of detergent resistant membrane fractions in which MPZ and PMP22 can partition in biophysical experiments ^29 73^. Myelin sheets themselves are high in cholesterol ^92^ and PMP22 is cholesterol-phillic ^29,38,39,72,73^ and potentiates the organization of lipid raft proteins in myelin ^37^. In polarized epithelial cells, PMP22 localizes to tight junctions which are also organized by lipid ‘raft’ subdomains ^40 93^. Moreover, PMP22 is a claudin family member, with the closest relative Claudin-15 that forms extensive polymers within membranes to organize tight junctions ^83,94^. Thus, one possibility is that PMP22 associates with MPZ to help confine the resulting adhesion complex in microdomains allowing it to optimize its activity in the membrane. To test whether the PMP22~MPZ complex localized to plasma membrane subdomains, we performed immunogold labeling of freeze fracture replicas from HEK293 cells expressing MPZ-mCherry and PMP22-GFP (Figure 7) ^95^. Double gold labeling of MPZ and PMP22 revealed that both proteins were randomly distributed across the plasma membrane without an obvious concentration of proteins in specific plasma membrane subdomains. This included areas where the plasma membrane of one cell was in contact with another in case the formation of MPZ~MPZ interactions in *trans* between cells could form a more stable concentration of complexes. We did notice instances of small chains of intermembrane particles evident on the E-face (exofacial leaflet), but they were rare and we could not verify their identity by gold labeling, the vast majority of which was only evident on the P-face (cytosolic leaflet).

**Figure 6.**
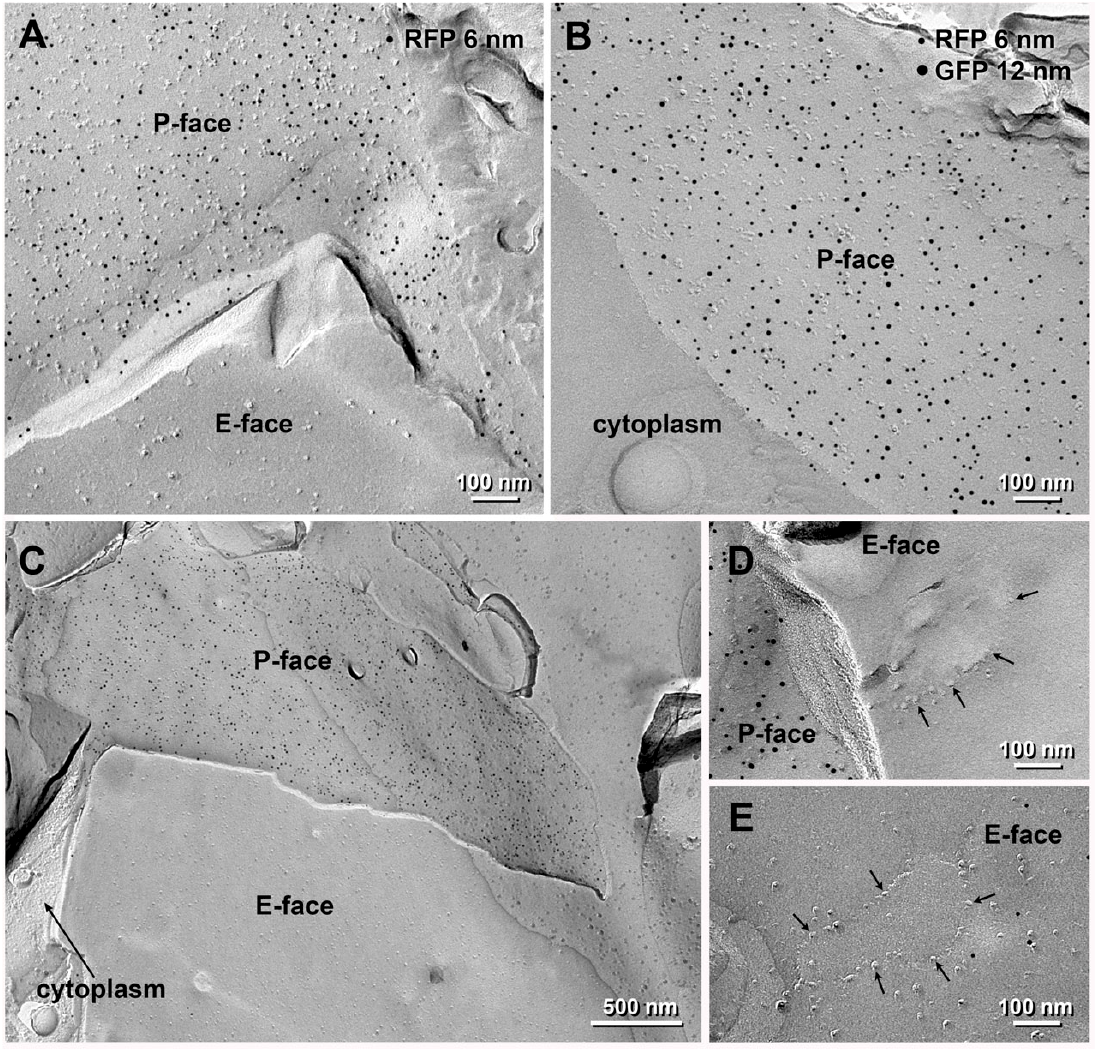
Immunogold labeling of MPZ and PMP22 in HEK293 cell freeze fracture replicas. **A**. Electron micrographs of the replicas from HEK293 cells expressing PMP-GFP only (A) and **B-E**. Micrographs of HEK293 cells expressing both MPZ-mCherry and PMP22-GFP. Cells were allowed to form cell-cell contacts before fixation in 1% paraformaldehyde. MPZ-mCherry was labelled with 6 nm immunogold particles and PMP22-GFP was labeled with 12 nm immunogold particles. Where two cell membranes are contacting, the immuno-gold labeling for both proteins were observed exclusively on the protoplasmic face/leaflet (P-face) and always homogeneously spread throughout the plasma membrane rather than distributed in clusters. The labeling patter was same in the single expression and labeling (**A**) and double expression and labeling (**B**). The topology of two contacting membrane is presentenced in (**C**). The labeled P-face shows continuity to its cytoplasm and an extraplasmic face/leaflet (E-face) of another cell is closely contacting on the P-face. The P-face exclusive labeling indicates the specificity of the labeling and consistent with the intercellular epitope locations. There were instances of intramembrane particles in a small cluster and strings observed on the E-face (**D, E**, arrows), but these were only rarely found. The small amount of immunogold particles near the string in (**E**) indicated the labeling of a hidden P-face below the E-face, called cryptic labeling, however the positions of the immunogold particles didn’t follow the string shape and could not be verified if they correspond to the localization of PMP22-GFP or MPZ-mCherry.

## Discussion

Here we find that two major myelin proteins, PMP22 and MPZ, form a strong and specific complex that is mediated by interactions between their transmembrane domains. That PMP22 and MPZ can form a complex has been previously proposed along with the model that their interaction is mediated by their extracellular domains potentially working in *trans* to bridge myelin wraps ^8,50,67^. Those studies report using the non-ionic detergents NP-40 and Triton X-100 with which we were unable to recover any associated MPZ in PMP22 co-immunoprecipitates. Rather the integrity of the complex was preserved using DDM and CHS in which MPZ very efficiently associated with PMP22. Despite the paucity of insight into PMP22 function and the potential importance of the PMP22~MPZ complex, analysis of a PMP22~MPZ complex has not been pursued by other groups in the intervening 2 decades. We suggest that the remarkable dependence on particular detergent conditions described here for the preservation of the PMP22~MPZ complex may have been an impediment for further study. Previous data also indicated that MPZ may form its interaction with PMP22 *via* its extracellular Ig-like domain. However, these studies used a recombinant GST-MPZ-Ig fusion protein produced intracellularly in bacteria. The MPZ Ig-like domain maintains its structure *via* an intradomain disulfide bond which is perturbed in the reduced bacterial cytoplasmic environment, leading to production of a mis-folded aggregate ^22^. Similarly, we found that intracellularly produced recombinant MPZ Ig-like domain fused to GST, the latter also containing a reactive cysteine, also produces a misfolded aggregate. Use of a structurally impaired Ig domain undermines the ability to make conclusions about how the MPZ Ig domain may or may not interact with partner proteins. In contrast, our data support the model that PMP22 and MPZ interact in the same membrane bilayer *via* their TMDs. This is supported by chimeric proteins made of MPZ and its related single Ig-like domain family members whereby proteins with the MPZ TMD were able to interact with PMP22, whereas substitution of the extracellular domain of MPZ with that of SCN2B or MPZL1 had no effect on interaction with PMP22. This model was further verified with amino acid substitutions in both MPZ and PMP22 that perturb complex formation and cluster within the TMD of MPZ and the 2^nd^ transmembrane helix of PMP22, respectively. In addition, we find that the PMP22~MPZ complex can be recovered when both are expressed in the same cell, but not when each is expressed in separate cell populations and mixed after cell lysis, and that interaction determined by FRET occurred within the plasma membrane segments that were not juxtaposed to other cells. These data further rationalize the model in Figure 5 that the PMP22~MPZ forms in *cis* within the same membrane.

PMP22 plays a crucial role in myelin formation and its homozygous loss results in severe demyelinating neuropathy ^62,96,97^. Heterozygous loss of PMP22 results in a milder peripheral neuropathy, HNPP (Hereditary Neuropathy with Pressure Palsies). We found the A67T patient variant of PMP22 that also causes a partial loss-of-function HNPP-like phenotype also blocks MPZ association but preserves PMP22 trafficking and interactions with other proteins interrogated here. These data strongly suggest that association of PMP22 with MPZ is an important requirement for PMP22 to function in myelin. This poses two complementary questions about the molecular and cellular functions of the PMP22~MPZ complex.

One question is what the role of MPZ~PMP22 complex formation is to promote normal myelination. Overall, little is known about what PMP22 does at the molecular level. PMP22 binds cholesterol and can distort lipid bilayers, organize lipid subdomains, coordinate the actin cytoskeleton dynamics, and in some cells coordinate formation of cell-cell tight junctions ^33,35,37-40,73^. PMP22 is also required for proper biogenesis of junctions that maintain the morphology of the paranodal loops of myelin ^52^. MPZ is largely restricted to compact myelin that is enriched in cholesterol and proper levels of cholesterol are required for the trafficking of MPZ to myelin wraps in compact myelin ^98,99^. One possibility is that PMP22 helps organize lipid subdomains that cluster MPZ to better fulfill its role as an adhesion protein. Alternatively, PMP22 itself might directly perform adhesive activity that amplifies the adhesion properties of MPZ to which it is complexed. PMP22 and MPZ segregate to detergent-resistant membranes, indicating a potential for populating lipid rafts ^29-31^ and X-ray diffraction data from myelin is consistent with the presence of dynamic yet small discrete membrane subdomains within condensed myelin ^27^. Freeze fracture data of myelin show the presence of larger intramembrane particles within both the P and E-faces that may correspond to multiple MPZ proteins and potentially could have PMP22 present as well to organize MPZ tetramers ^24-26^. Moreover, current models for how MPZ is organized across the intraperiod line propose lateral networks of proteins that PMP22 association could potentially help configure ^17,20^. In our freeze fracture experiments, we did not observe a focused concentration of MPZ or PMP22 in the plasma membrane even in areas where cell-to-cell contacts were present and potentially allowing MPZ-mediated adhesion. We also did not observe the type of intramembrane particles that were observed in myelin or in cells expressing high levels of claudin-15, a close structural homolog of PMP22 that forms large tight junction arrays. Our data in HEK293 cells do not exclude these possibilities may yet happen in myelin, and may rather indicate that other factors in myelin are required to achieve such structures. This could involve the very high levels of MPZ and PMP22 in myelin membranes, the high level of cholesterol in myelin membranes, and the confined space these proteins operate in as exemplified by the very thin cytoplasmic space that forms the major dense line.

There are other possibilities for what functions are fulfilled when PMP22 and MPZ form a complex. There may also be effects on the biogenesis of PMP22 that are aided by association with MPZ. Complex formation could potentially control the levels and distribution of PMP22. A sizable fraction of PMP22 is degraded in Schwann cells rather than being sent to the plasma membrane ^63^. In heterologous cells, a large proportion of PMP22 is trapped in the ER and that proportion increases with increased expression of PMP22 ^65^. PMP22 can also be found in aggresomes, suggesting that PMP22 is inefficiently degraded by ER-associated degradation (ERAD) ^36,100^. Complex formation with MPZ might potentiate the ability of PMP22 to exit the ER or help control the levels of PMP22 by regulating its recognition by the ERAD machinery.

A second question is whether complex formation with MPZ plays a pathogenic role in CMT, especially CMT1A that is linked to a gene duplication of *PMP22* resulting in disease caused by mild over-expression of wildtype PMP22. One potential mechanism for the pathogenesis of CMT1A may be due to ER-stress from excess production of PMP22 ^61,64,65,101^, that could be exacerbated by alteration of membrane morphology and perturbation of cholesterol distribution ^38,39,49^. These effects would be mediated by simply too much PMP22 in the ER and motivates the therapeutic strategy of reducing PMP22 levels ^102^. However, CMT1A could be caused not simply by more PMP22 *per se*, but rather too much of the PMP22~MPZ complex. Alternatively, the ability of excess PMP22 to form a complex with MPZ may be protective, reducing the severity of CMT that excess PMP22 would otherwise cause if it were unable to form a complex with MPZ. Either of these possibilities would open new therapeutic opportunities to find drugs that could weaken or strengthen the association of PMP22 with MPZ. Now that we have defined amino acid substitutions in MPZ and PMP22 that specifically block complex formation but preserve other protein interactions and proper trafficking to the cell surface, experiments aimed at understanding the cellular and physiological functions of the PMP22~MPZ complex *in vivo* and its potential role in driving pathogenesis of CMT1A can now be addressed.

A full explanation of the normal function and pathogenic functions will need to incorporate the fact that PMP22 interacts with additional partners. PMP22 associates with itself to form dimers and also interacts with Jam-C and myelin-associated glycoprotein (MAG) ^52,54,55^. Here we show additional interactions with Jam-B (Jam2) and EMP1 (Figure.6). Mutant forms of PMP22 trapped in the ER can also form a complex with STIM1, a subunit of the store-operated calcium channel ^35^. These interactions could be critical for these PMP22 partners to operate. JAM-C is expressed in junctional regions of Schwann cells and MAG is localized to the myelin wrap closest to the axon ^103,104^. Loss of Jam-C or MAG result in neuropathy similar to loss of PMP22 ^52^. EMP1 expression is stimulated upon nerve injury and is inversely regulated compared to PMP22, which is induced during Schwann cell differentiation ^105^. An important question is how PMP22 is properly portioned among these partners and how that portioning might be influenced by more or less PMP22, both conditions of which are pathogenic. Thus, it may not be the loss of MPZ association *per se* that is responsible for the HNPP phenotype in the PMP22 A67T variant, but an imbalance in other PMP22 complexes. The many potential roles and partners for PMP22 also advance the question of the overall stoichiometry of these complexes and whether there is ‘enough’ PMP22 to associate with all these partners. Previous experiments that estimated the relative abundance of MPZ and PMP22 indicated that while ~50% of myelin protein is MPZ, only 5% is PMP22 ^7,106-108^. However, these estimates come from dyeing proteins after SDS-PAGE, which does not accurately compare proteins with very different amino acid content. Moreover, PMP22 abundance, in terms of mRNA levels, has been shown to be twice that of MPZ ^109^. Thus, there may be a larger ratio of PMP22 to MPZ and its binding partners than we currently realize. Nonetheless, even these upper estimates are unlikely to allow all of MPZ and other partners to be in a complex with PMP22. Thus, it may be that PMP22 plays a more acute transitory role in the function of its binding partners.

### Limitations of this study

Here we show how MPZ can form a complex with PMP22 and show that a PMP22 variant associated with a loss-of-function phenotype in people specifically blocks formation of the complex. The implication is that complex formation is critical for PMP22 to achieve its full function. However, this is one particular patient variant and we cannot know if is some other PMP22 propertiy affected by that variant that might explain loss-of-function. Using the structural insights we obtained through mutagenesis, we have identified other mutations that cause loss of MPZ~PMP22 association without disturbing the other cellular and biochemical behaviors and interactions that we do know about. These should be tested *in vivo* to extend the structure/function correlation to better test the importance of the MPZ~PMP22 interaction and allow for a detailed analysis of what this complex achieves at the cellular and molecular level. We also note that all of our experiments are in heterologous cells. We did not investigate how abundant this complex is in normal myelin nor where those interactions occur within myelin. The heterologous cells also did not recapitulate the types of membrane protein structures in myelin that previous studies speculated were comprised of MPZ, so we cannot assess whether PMP22 association with MPZ is an important structural component of the adhesion structures thought to operate between authentic myelin sheets across the intraperiod line. Nonetheless, the mutations defined here that disrupt the complex would enable these types of *in vivo* studies. Finally, we did not explore how these proteins might make higher order oligomers. We could detect MPZ~MPZ, PMP22~PMP22, and MPZ~PMP22 interactions but these proteins did not form larger polymers in our analysis. Our data neither rule-in nor rule-out that such assemblies can form and whether they operate within myelin.

## Acknowledgements

We acknowledge the University of Iowa personnel and instrumentation in the IIHG Genomic Sequencing, the Carver College of Medicine NMR, and Protein & Crystallography core facilities, supported by the Roy J. and Lucille A. Carver College of Medicine and grants from the Roy J. Carver Charitable Trust. We thank Miranda Schene and Sandipan Chowdhury for help with detergent and FPLC analysis. This work was supported by NIH-R01 R35GM148239 to CAA, U54NS065712 to MES, and NIH RO1GM058202 to RCP. CAA and RCP were supported by the Roy J. Carver Charitable Trust. This paper was typeset with the bioRxiv word template by @Chrelli: www.github.com/chrelli/bioRxiv-word-template

## Competing interest statement

The authors have no competing interests to declare.

## Materials and Methods

### Plasmids, antibodies, cell culture, and materials

Gene synthesized open reading frames were purchased from IDT (Coralville, IA), subcloned into expression plasmids by Gibson assembly (e.g. pcDNA3.1, ThermoFisher, Waltham, MA), and sequenced by Sanger sequencing and/or Oxford nanopore long-read technology using Plasmidsaurus (Eugene, OR). Plasmid use and description are provided in *Supplementary Table 1* and the full sequence of plasmids provided in additional supplementary materials. GFP tagging of MPZ, PMP22, and Glycophorin A was done using msGFP2 (Valbuena et al., 2020), a monomeric superfolder version of GFP that was stable against photobleaching. Antibodies used were: mouse α-GFP monoclonal (sc-9996, Santa Cruz Technology, Santa Cruz, CA), mouse α-HA monoclonal (901514, BioLegend, San Diego, CA), mouse α-myc-Tag (9B11) monoclonal (2276, Cell Signaling Technology, Danvers, MA), rabbit α-mCherry polyclonal (AB356482, EMD Millipore Corp., Burlington, MA). Peroxidase-conjugated (HRP) α-mouse (cat#:70776S) or α-rabbit secondary antibody (cat# 7074S) were obtained from Cell Signaling Technology. Dodecyl-β-D-maltoside (DDM) and cholesteryl hemisuccinate (CHS) from Anatrace (Maumee, OH) were kept as a stock solution (10% and 2%, respectively) in 200mM Tris pH 8.0 at 4°C. Protease inhibitors cOmplete lacking EDTA, and Pefabloc, were obtained from Sigma (St. Louis, MO).

HEK293 cells (ATCC: CRL-1573) and RT4-D6P2T Schwannoma cells (ATCC: CRL-2768) were cultured at 37°C with 5% CO2 in Dulbecco’s modified Eagle’s medium (DMEM) supplemented with 10% fetal bovine serum (FBS) and 1% penicillin + streptomycin antibiotics (Gibco, Grand Island, NY). Transient transfections were accomplished with Lipofectamine LTX using 5 µg plasmid DNA per million cells in Opti-MEM reduced serum media according to manufacturer’s instruction (ThermoFisher, Waltham, MA). After transfection, cells were cultured in DMEM with 10% FBS without antibiotics prior to analysis. Stable transfected PMP22-myc^exo^ cell lines were made using the piggyBac system (Ding et al., 2005) and a co-expression vector with the puromycin-resistance protein fused to blue fluorescent protein (mTAGBFP2) appended with a nuclear localization signal. At 24 hrs post transfection, cells were grown in DMEM media with 10% FBS and 5 µg/ml puromycin (Sigma, St. Louis, MO). Cells expressing mTAGBFP2 were sorted from non-mTAGBFP2 expressing cells by flow cytometry.

### Protein expression in cultured cells

Transient transfections were accomplished with Lipofectamine LTX using 5 µg plasmid DNA per million cells in Opti-MEM reduced serum media according to manufacturer’s instruction (ThermoFisher, Waltham, MA). After transfection, cells were cultured in DMEM with 10% FBS without antibiotics prior to analysis. Stable transfected PMP22-myc^exo^ cell lines were made using the piggyBac system and a co-expression vector with the puromycin-resistance protein fused to blue fluorescent protein (mTAGBFP2) appended with a nuclear localization signal ^110,111^. At 24 hrs post transfection, cells were grown in DMEM media with 10% FBS and 5 µg/ml puromycin (Sigma, St. Louis, MO) for 2 days. Cells expressing mTAGBFP2 were sorted from non-mTAGBFP2 expressing cells by flow cytometry and cultured in the absence of puromycin.

### Immunoprecipitation

Anti-GFP nanobody-conjugated agarose beads (Protein Crystallograpy facility, University of Iowa) were used for immunoprecipitations. Cells were grown in 6 cm dishes, 18-20 hrs post transfection washed with Dulbecco’s phosphate-buffered saline (DPBS) and suspended with Versene (ThermoFisher, Waltham, MA). Cells were lysed for 30 min on ice in 500 µl lysis buffer (PBS with 1% DDM, 0.2% CHS, supplemented with cOmpleteTM, and Pefabloc^®^ protease inhibitors) and centrifuged at 4°C 12,200 rpm for 15 min. The cleared supernatant (400 µl) was mixed with 34 µl of α-GFP nanobody agarose beads in 100 µl PBS and incubated 1 hr rotating at 4°C. After three washes by centrifugation at 5,000 rpm 4 min with 1ml of cold lysis buffer diluted 8 times with PBS, proteins were eluted with SDS-loading sample buffer with heating at 95°C. Equal amounts of whole cell lysates (input) and immunoprecipitated proteins were resolved by SDS-PAGE then immunoblotted with indicated antibodies.

### Fluorescence microscopy

HEK293 cells were grown on glass bottom dishes (MatTek, P35G-1.5-20-C). After 18-20 hrs of transfection, live cells expressing either GFP and/or mCherry tagged proteins were visualized in the culture media using a Leica SP8 confocal microscope equipped with 405, 488 and 552 nm lasers.

For FRET analysis, cells were fixed in 4% paraformaldehyde in DPBS, washed with 50 mM Tris-HCl in DPBS pH7.5 to inactivate residual fixative, and imaged in DPBS. The Leica LAX X Microlab software was used using the FRET AB acceptor photobleaching method with the msGFP/mCherry pair of fluorophores ^112^. For visualization of stable cell lines with wild type or mutant PMP22-myc^exo^ at the cell surface, cells were grown on glass coverslips in 6-well culture plates, fixed with 4% paraformaldehyde in DPBS 15 min, washed with 50 mM Tris-HCl in DPBS pH 7.5, then washed with DPBS. Coverslips were blocked with 5% goat serum in DPBS for 30 min, then incubated with mouse α-myc-tag antibody 40 min at RT. Following three washes with DPBS, coverslips were incubated with goat α-mouse Alexa568 secondary antibody for 30 min in the dark. Following three washes with DPBS, the coverslips were mounted on glass slides using Vectashield antifade mounting medium (H-1000, Vector Laboratories, Burlingame, CA).

### Flow Cytometry

HEK293 cells were lifted from plates using trypsin/EDTA, pelleted, and resuspended in 5 mls of media (Dulbecco’s modified Eagle’s medium (DMEM) supplemented with 10% fetal bovine serum (FBS) and 1% penicillin + streptomycin antibiotics). Cells were filtered through 40µM nylon mesh (Thermofisher, Waltham, MA), and stored for up to 1 hr at 4°C prior to Flow analysis and/or cell sorting. Flow cytometry data were obtained using a Becton Dickinson LSRII equipped with 405 nm, 488 nm, and 561 nm lasers. Analysis and data graphing was done using FlowJo v10 software (FlowJo, Ashland, OR 97520). To isolate BFP positive HEK293 cells after integration of piggyBac-based vectors expressing nuclear localized BFP, suspended and filtered cells were sorted for BFP using a Becton Dickinson Aria Fusion equipped with 407 nm, 488 nm, and 639 nm lasers. The gating strategy used FSC and SSC gates to define the median range of cell size and quality. For cell sorting, untransfected HEK293 cells were used for calibration such that the gate BFP+ cells allowed <1% of untransfected HEK293 cells to pass. No upper limit on BFP expression was set for this gate.

### Size exclusion chromatography

HEK293 cells expressing PMP22-GFP alone or with MPZ-HA were lysed in PBS with 1% DDM, 0.2% CHS buffer, supplemented with cOmpleteTM, and Pefabloc^®^ protease inhibitors and centrifuged at 4°C 12,200 rpm for 15 min, then at 100,000xg for 30 min. Solubilized lysate sample was separated on Superose 6 10/300 column (Cytiva) by LC (HPLC, Shimadzu or NGC Chromatography System, BioRad) equilibrated with PBS buffer pH 7.4 containing 0.05% DDM and 0.01% CHS, at 0.5 mL/min, and observed by fluorescence detection of the GFP fusion protein (Ex/Em: 488/510 nm) and/or absorbance at A280. With the BioRad System, 1 mL fractions were collected for further analysis.

### Electron Microscopy

Transiently transfected HEK293 cells grown to confluence were fixed by removing half the volume of the culture media and replacing it with 2% PFA in 0.12M Sorensen’s phosphate buffer (PB) for 15 min. The fixative was replaced with fresh 2% PFA in 0.12M PB and incubated for 1 hr at room temperature. Fixed cells were scrapped, pelleted and processed for freeze fracture replica immunogold labeling as previously described in ^95,113^. Briefly, the pellets of cells were cryoprotected in 10%, 20%, 30% glycerol in 0.1M PB respectively for one hr each, and stored in 30% glycerol in 0.1M PB for overnight. A drop of the cell pellet was placed on a cylinder-type carrier (1.0 mm cat# 16770134, Leica) and covered with a dome-type carrier (0.8 mm cat#16770132, Leica) coated with a lecithin, then high-pressure frozen with an HPM 100 (Leica Microsystems Inc., Deerfield, IL). The frozen cells were knife-fractured at −120°C, and replicated with a 2 nm pre-carbon deposition, shadowed with 3 nm of carbon/platinum at a 60° angle, and supported by a final deposition of 30-40 nm of carbon using a JFDII Freeze Fracture machine (JEOL, Akishima, Japan). The replicas were digested in a solution containing 2.5% SDS, 20% sucrose and 15 mM Tris-HCl (pH 8.3) with a gentle agitation at 82.5°C for 19 hrs. The replicas were washed and blocked with 4% bovine serum albumin and 1% fish skin gelatin in 50 mM Tris buffered saline (TBS, pH 7.6) for 1 hr, then incubated with rabbit anti-RFP(mCherry) antibody (final concentration of 1.1mg/ml) and chicken anti-GFP (final concentration of 0.25µg/ml) diluted in the same blocking solution for 18 hours at room temperature. After several washes, the replicas were incubated in a solution containing donkey anti-rabbit IgG conjugated to 6 nm gold particles (1:30, Jackson Immunoresearch) and donkey anti-chicken IgY++ (IgG) conjugated to 12 nm gold particles (1:30 Jackson Immunoresearch) for 18 hrs at room temperature. After being washed, the replicas were picked up on 2 mm aperture copper grids and examined with a Tecnai G2 Spirit Bio twin transmission electron microscope (ThermoFisher, Waltham, MA) at 100 kv acceleration voltage. Images were taken with a Veleta CCD camera (Olympus, Tokyo, Japan) controlled by a TIA software (ThermoFisher, Waltham, MA, formerly FEI, Hillsboro, OR). Brightness and contrast of images were maximumly adjusted in Photoshop (Adobe CS6).

### Modeling

Models for PMP22 (AF-Q01453-F1) and MPZ (AF-P25189-F1) were obtained from the AlphaFold PSDB ^84^. The complex between PMP22 (all residues) and MPZ (transmembrane residues: R153-L184) was modelled using the HADDOCK webserver ^114,115^ with initial restraints defined by all residues shown to disrupt the complex when experimentally mutated (on PMP22: A67, I70, I71, I73, I74, F75, I116; on MPZ: G155, V157, L158, I162, L166, V169). The best models from the two top ranking MPZ docking poses exhibited mutually exclusive interactions with restraints and were each further refined on a subset of residue restraints (Figure S1). The final model that best rationalizes the mutagenesis data was constrained for involvement of I70 without a requirement for proximity of I116. Full-length MPZ (residues I30-K248) was superimposed ^116^ on the final model and embedded in a representative lipid bilayer (60%POPC:40%cholesterol) using the CHARMM-GUI webserver ^117^ with membrane orientation optimized by the PPM method ^118^.

### Statistical analysis

FRET data as well as colocalization data were graphed using Prism software (Graphpad Software, Boston, MA 02110). One-way ANOVA analysis of data with Tukey’s multiple comparison tests were made using Prism. To measure co-localization, the JaCoP plugin ^119^ for Fiji was used to find the Manders coefficient ^120^.

## Key Resources Table

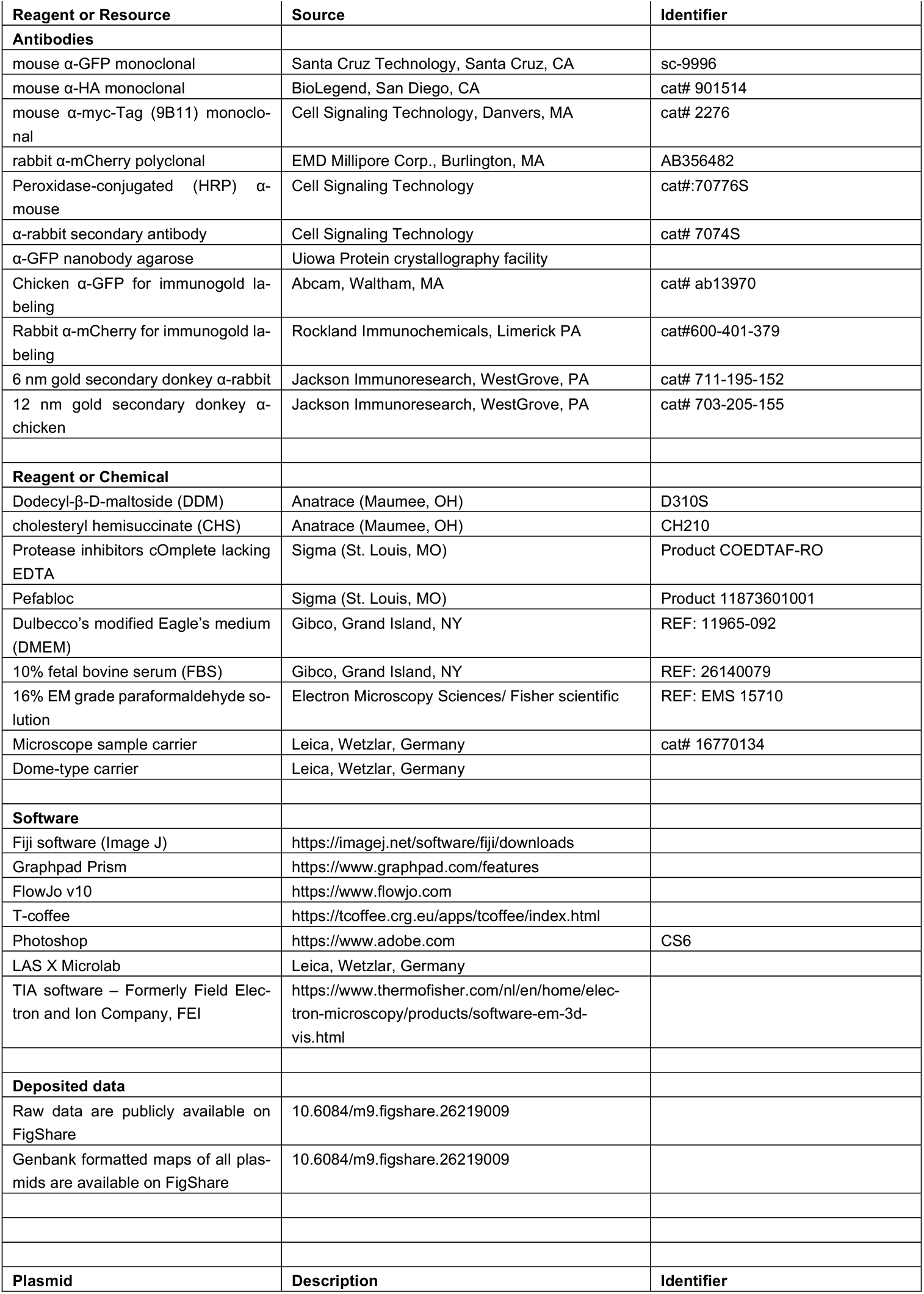
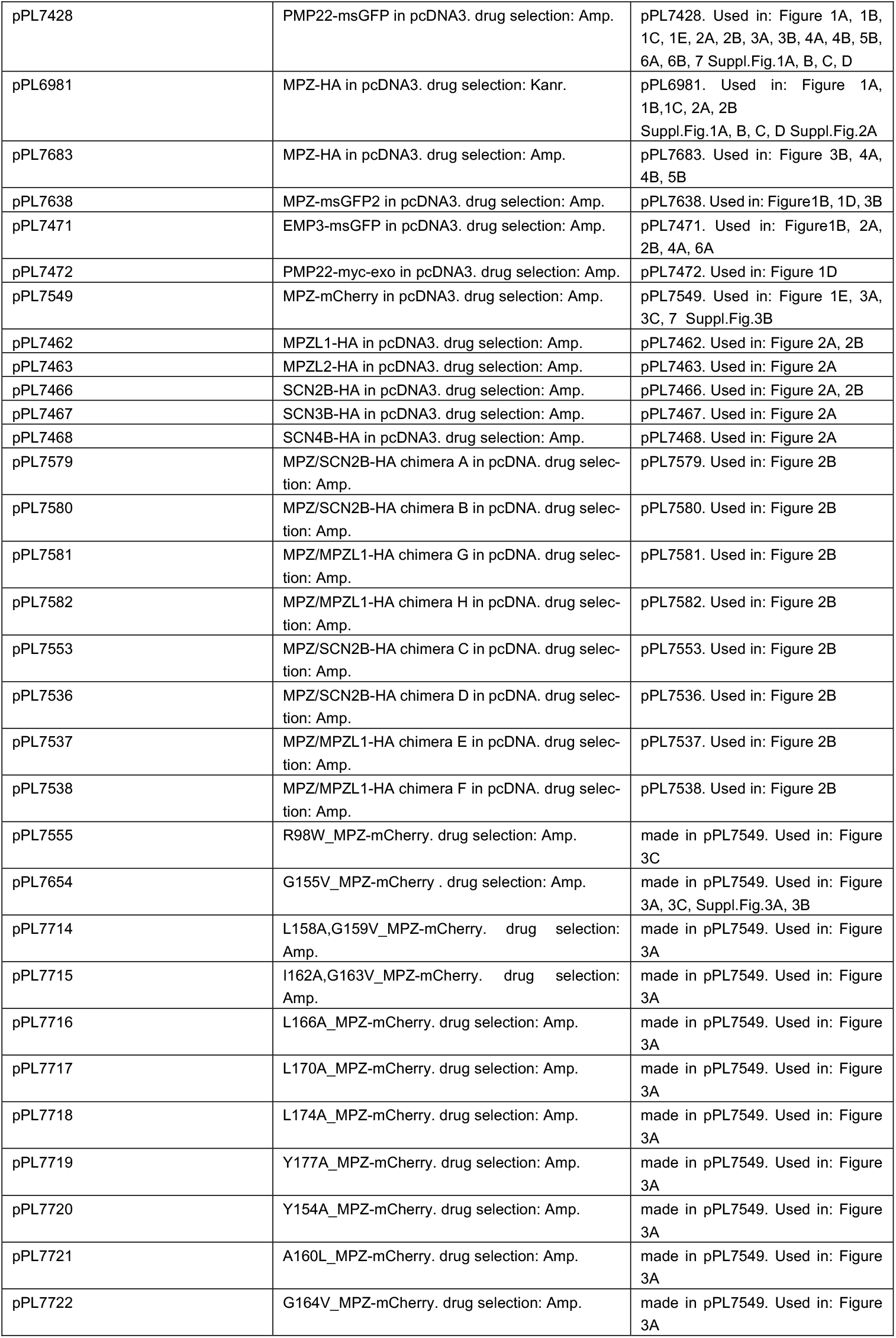
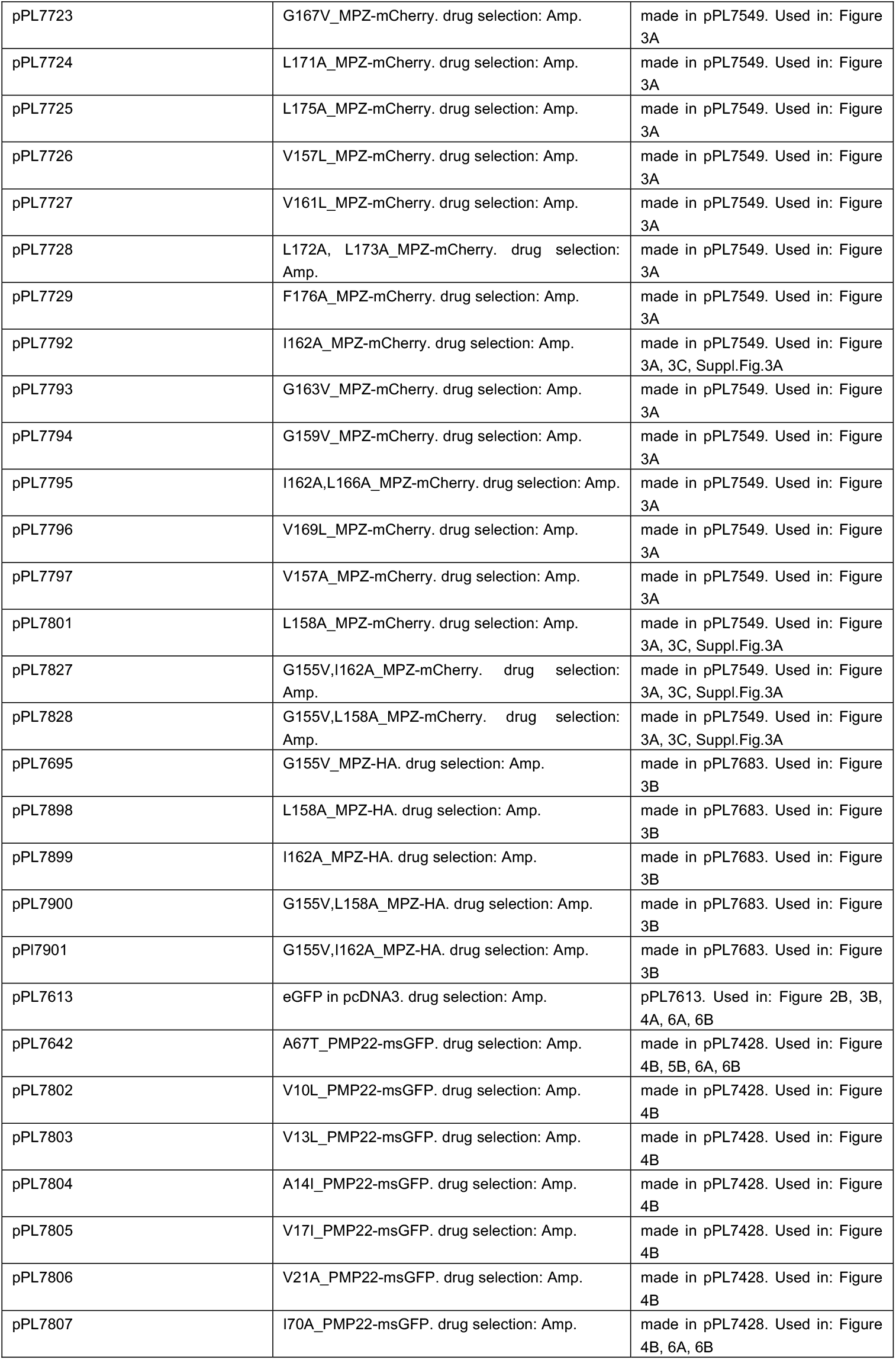
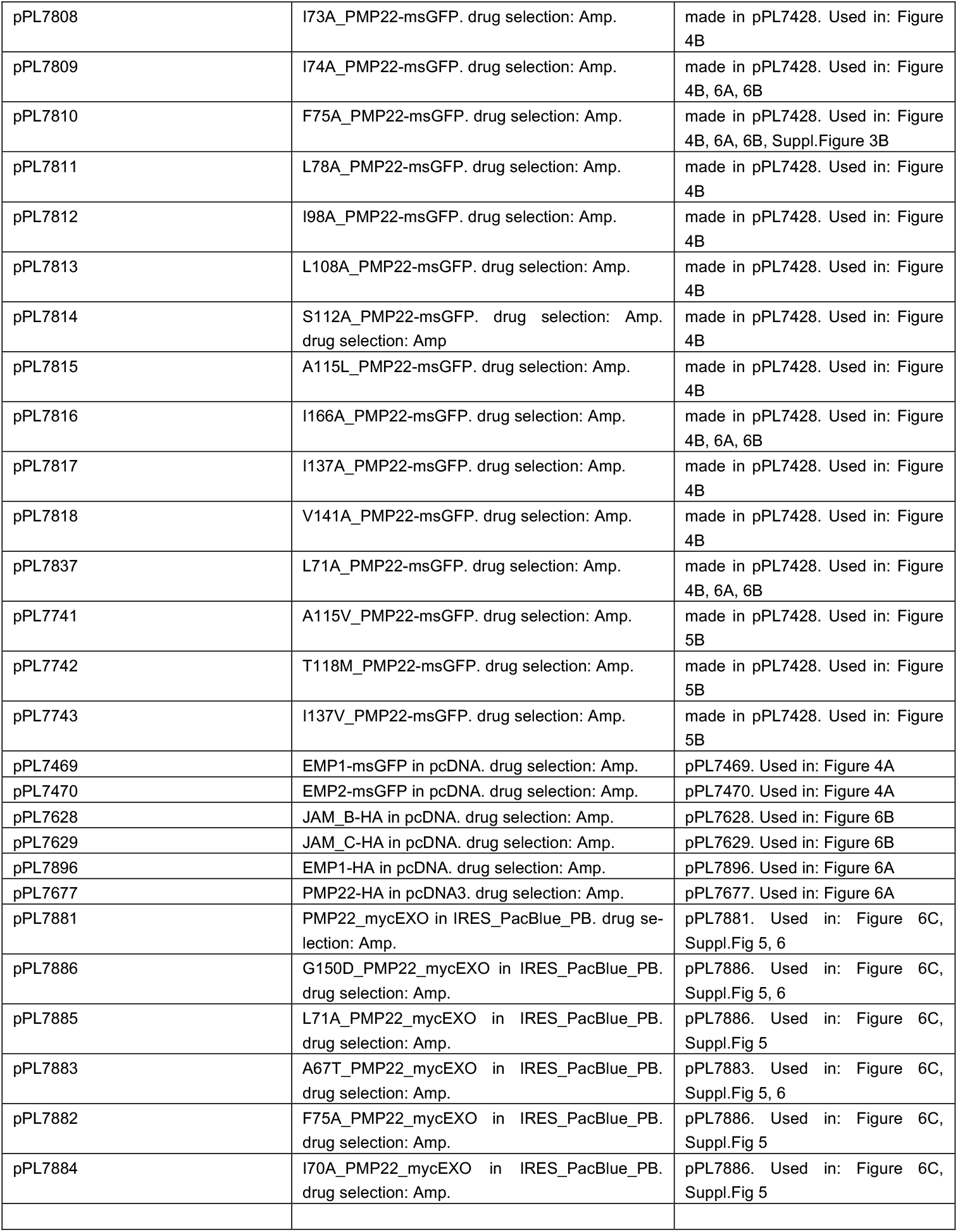

| Reagent or Resource | Source | Identifier |
| --- | --- | --- |
| **Antibodies** |  |  |
| mouse α-GFP monoclonal | Santa Cruz Technology, Santa Cruz, CA | sc-9996 |
| mouse α-HA monoclonal | BioLegend, San Diego, CA | cat# 901514 |
| mouse α-myc-Tag (9B11) monoclonal | Cell Signaling Technology, Danvers, MA | cat# 2276 |
| rabbit α-mCherry polyclonal | EMD Millipore Corp., Burlington, MA | AB356482 |
| Peroxidase-conjugated (HRP) α-mouse | Cell Signaling Technology | cat#:70776S |
| α-rabbit secondary antibody | Cell Signaling Technology | cat# 7074S |
| α-GFP nanobody agarose | Uiowa Protein crystallography facility | |
| Chicken α-GFP for immunogold labeling | Abcam, Waltham, MA | cat# ab13970 |
| Rabbit α-mCherry for immunogold labeling | Rockland Immunochemicals, Limerick PA | cat#600-401-379 |
| 6 nm gold secondary donkey α-rabbit | Jackson ImmunoResearch, WestGrove, PA | cat# 711-195-152 |
| 12 nm gold secondary donkey α-chicken | Jackson ImmunoResearch, WestGrove, PA | cat# 703-205-155 |
| **Reagent or Chemical** |  |  |
| Dodecyl-β-D-maltoside (DDM) | Anatrace (Maumee, OH) | D310S |
| cholesteryl hemisuccinate (CHS) | Anatrace (Maumee, OH) | CH210 |
| Protease inhibitors cOmplete lacking EDTA | Sigma (St. Louis, MO) | Product COEDTAF-RO |
| Pefabloc | Sigma (St. Louis, MO) | Product 11873601001 |
| Dulbecco's modified Eagle's medium (DMEM) | Gibco, Grand Island, NY | REF: 11965-092 |
| 10% fetal bovine serum (FBS) | Gibco, Grand Island, NY | REF: 26140079 |
| 16% EM grade paraformaldehyde solution | Electron Microscopy Sciences/ Fisher scientific | REF: EMS 15710 |
| Microscope sample carrier | Leica, Wetzlar, Germany | cat# 16770134 |
| Dome-type carrier | Leica, Wetzlar, Germany |  |
| **Software** |  |  |
| Fiji software (Image J) | https://imagej.net/software/fiji/downloads |  |
| Graphpad Prism | https://www.graphpad.com/features |  |
| FlowJo v10 | https://www.flowjo.com |  |
| T-coffee | https://tcoffee.crg.eu/apps/tcoffee/index.html |  |
| Photoshop | https://www.adobe.com | CS6 |
| LAS X Microlab | Leica, Wetzlar, Germany |  |
| TIA software – Formerly Field Electron and Ion Company, FEI | https://www.thermofisher.com/nl/en/home/electron-microscopy/products/software-em-3d-vis.html |  |
| **Deposited data** |  |  |
| Raw data are publicly available on FigShare | 10.6084/m9.figshare.26219009 |  |
| Genbank formatted maps of all plasmids are available on FigShare | 10.6084/m9.figshare.26219009 |  |
| **Plasmid** | **Description** | **Identifier** |
| pPL7428 | PMP22-msGFP in pcDNA3. drug selection: Amp. | pPL7428. Used in: Figure 1A, 1B, 1C, 1E, 2A, 2B, 3A, 3B, 4A, 4B, 5B, 6A, 6B, 7 Suppl.Fig.1A, B, C, D |
| pPL6981 | MPZ-HA in pcDNA3. drug selection: Kanr. | pPL6981. Used in: Figure 1A, 1B, 1C, 2A, 2B<br>Suppl.Fig.1A, B, C, D Suppl.Fig.2A |
| pPL7683 | MPZ-HA in pcDNA3. drug selection: Amp. | pPL7683. Used in: Figure 3B, 4A, 4B, 5B |
| pPL7638 | MPZ-msGFP2 in pcDNA3. drug selection: Amp. | pPL7638. Used in: Figure1B, 1D, 3B |
| pPL7471 | EMP3-msGFP in pcDNA3. drug selection: Amp. | pPL7471. Used in: Figure1B, 2A, 2B, 4A, 6A |
| pPL7472 | PMP22-myc-exo in pcDNA3. drug selection: Amp. | pPL7472. Used in: Figure 1D |
| pPL7549 | MPZ-mCherry in pcDNA3. drug selection: Amp. | pPL7549. Used in: Figure 1E, 3A, 3C, 7 Suppl.Fig.3B |
| pPL7462 | MPZL1-HA in pcDNA3. drug selection: Amp. | pPL7462. Used in: Figure 2A, 2B |
| pPL7463 | MPZL2-HA in pcDNA3. drug selection: Amp. | pPL7463. Used in: Figure 2A |
| pPL7466 | SCN2B-HA in pcDNA3. drug selection: Amp. | pPL7466. Used in: Figure 2A, 2B |
| pPL7467 | SCN3B-HA in pcDNA3. drug selection: Amp. | pPL7467. Used in: Figure 2A |
| pPL7468 | SCN4B-HA in pcDNA3. drug selection: Amp. | pPL7468. Used in: Figure 2A |
| pPL7579 | MPZ/SCN2B-HA chimera A in pcDNA. drug selection: Amp. | pPL7579. Used in: Figure 2B |
| pPL7580 | MPZ/SCN2B-HA chimera B in pcDNA. drug selection: Amp. | pPL7580. Used in: Figure 2B |
| pPL7581 | MPZ/MPZL1-HA chimera G in pcDNA. drug selection: Amp. | pPL7581. Used in: Figure 2B |
| pPL7582 | MPZ/MPZL1-HA chimera H in pcDNA. drug selection: Amp. | pPL7582. Used in: Figure 2B |
| pPL7553 | MPZ/SCN2B-HA chimera C in pcDNA. drug selection: Amp. | pPL7553. Used in: Figure 2B |
| pPL7536 | MPZ/SCN2B-HA chimera D in pcDNA. drug selection: Amp. | pPL7536. Used in: Figure 2B |
| pPL7537 | MPZ/MPZL1-HA chimera E in pcDNA. drug selection: Amp. | pPL7537. Used in: Figure 2B |
| pPL7538 | MPZ/MPZL1-HA chimera F in pcDNA. drug selection: Amp. | pPL7538. Used in: Figure 2B |
| pPL7555 | R98W_MPZ-mCherry. drug selection: Amp. | made in pPL7549. Used in: Figure 3C |
| pPL7654 | G155V_MPZ-mCherry. drug selection: Amp. | made in pPL7549. Used in: Figure 3A, 3C, Suppl.Fig.3A, 3B |
| pPL7714 | L158A,G159V_MPZ-mCherry. drug selection: Amp. | made in pPL7549. Used in: Figure 3A |
| pPL7715 | I162A,G163V_MPZ-mCherry. drug selection: Amp. | made in pPL7549. Used in: Figure 3A |
| pPL7716 | L166A_MPZ-mCherry. drug selection: Amp. | made in pPL7549. Used in: Figure 3A |
| pPL7717 | L170A_MPZ-mCherry. drug selection: Amp. | made in pPL7549. Used in: Figure 3A |
| pPL7718 | L174A_MPZ-mCherry. drug selection: Amp. | made in pPL7549. Used in: Figure 3A |
| pPL7719 | Y177A_MPZ-mCherry. drug selection: Amp. | made in pPL7549. Used in: Figure 3A |
| pPL7720 | Y154A_MPZ-mCherry. drug selection: Amp. | made in pPL7549. Used in: Figure 3A |
| pPL7721 | A160L_MPZ-mCherry. drug selection: Amp. | made in pPL7549. Used in: Figure 3A |
| pPL7722 | G164V_MPZ-mCherry. drug selection: Amp. | made in pPL7549. Used in: Figure 3A |
| pPL7723 | G167V_MPZ-mCherry. drug selection: Amp. | made in pPL7549. Used in: Figure 3A |
| pPL7724 | L171A_MPZ-mCherry. drug selection: Amp. | made in pPL7549. Used in: Figure 3A |
| pPL7725 | L175A_MPZ-mCherry. drug selection: Amp. | made in pPL7549. Used in: Figure 3A |
| pPL7726 | V157L_MPZ-mCherry. drug selection: Amp. | made in pPL7549. Used in: Figure 3A |
| pPL7727 | V161L_MPZ-mCherry. drug selection: Amp. | made in pPL7549. Used in: Figure 3A |
| pPL7728 | L172A, L173A_MPZ-mCherry. drug selection: Amp. | made in pPL7549. Used in: Figure 3A |
| pPL7729 | F176A_MPZ-mCherry. drug selection: Amp. | made in pPL7549. Used in: Figure 3A |
| pPL7792 | I162A_MPZ-mCherry. drug selection: Amp. | made in pPL7549. Used in: Figure 3A, 3C, Suppl.Fig.3A |
| pPL7793 | G163V_MPZ-mCherry. drug selection: Amp. | made in pPL7549. Used in: Figure 3A |
| pPL7794 | G159V_MPZ-mCherry. drug selection: Amp. | made in pPL7549. Used in: Figure 3A |
| pPL7795 | I162A,L166A_MPZ-mCherry. drug selection: Amp. | made in pPL7549. Used in: Figure 3A |
| pPL7796 | V169L_MPZ-mCherry. drug selection: Amp. | made in pPL7549. Used in: Figure 3A |
| pPL7797 | V157A_MPZ-mCherry. drug selection: Amp. | made in pPL7549. Used in: Figure 3A |
| pPL7801 | L158A_MPZ-mCherry. drug selection: Amp. | made in pPL7549. Used in: Figure 3A, 3C, Suppl.Fig.3A |
| pPL7827 | G155V,I162A_MPZ-mCherry. drug selection: Amp. | made in pPL7549. Used in: Figure 3A, 3C, Suppl.Fig.3A |
| pPL7828 | G155V,L158A_MPZ-mCherry. drug selection: Amp. | made in pPL7549. Used in: Figure 3A, 3C, Suppl.Fig.3A |
| pPL7695 | G155V_MPZ-HA. drug selection: Amp. | made in pPL7683. Used in: Figure 3B |
| pPL7898 | L158A_MPZ-HA. drug selection: Amp. | made in pPL7683. Used in: Figure 3B |
| pPL7899 | I162A_MPZ-HA. drug selection: Amp. | made in pPL7683. Used in: Figure 3B |
| pPL7900 | G155V,L158A_MPZ-HA. drug selection: Amp. | made in pPL7683. Used in: Figure 3B |
| pPI7901 | G155V,I162A_MPZ-HA. drug selection: Amp. | made in pPL7683. Used in: Figure 3B |
| pPL7613 | eGFP in pcDNA3. drug selection: Amp. | pPL7613. Used in: Figure 2B, 3B, 4A, 6A, 6B |
| pPL7642 | A67T_PMP22-msGFP. drug selection: Amp. | made in pPL7428. Used in: Figure 4B, 5B, 6A, 6B |
| pPL7802 | V10L_PMP22-msGFP. drug selection: Amp. | made in pPL7428. Used in: Figure 4B |
| pPL7803 | V13L_PMP22-msGFP. drug selection: Amp. | made in pPL7428. Used in: Figure 4B |
| pPL7804 | A14I_PMP22-msGFP. drug selection: Amp. | made in pPL7428. Used in: Figure 4B |
| pPL7805 | V17I_PMP22-msGFP. drug selection: Amp. | made in pPL7428. Used in: Figure 4B |
| pPL7806 | V21A_PMP22-msGFP. drug selection: Amp. | made in pPL7428. Used in: Figure 4B |
| pPL7807 | I70A_PMP22-msGFP. drug selection: Amp. | made in pPL7428. Used in: Figure 4B, 6A, 6B |
| pPL7808 | I73A_PMP22-msGFP. drug selection: Amp. | made in pPL7428. Used in: Figure 4B |
| pPL7809 | I74A_PMP22-msGFP. drug selection: Amp. | made in pPL7428. Used in: Figure 4B, 6A, 6B |
| pPL7810 | F75A_PMP22-msGFP. drug selection: Amp. | made in pPL7428. Used in: Figure 4B, 6A, 6B, Suppl.Figure 3B |
| pPL7811 | L78A_PMP22-msGFP. drug selection: Amp. | made in pPL7428. Used in: Figure 4B |
| pPL7812 | I98A_PMP22-msGFP. drug selection: Amp. | made in pPL7428. Used in: Figure 4B |
| pPL7813 | L108A_PMP22-msGFP. drug selection: Amp. | made in pPL7428. Used in: Figure 4B |
| pPL7814 | S112A_PMP22-msGFP. drug selection: Amp.<br>drug selection: Amp | made in pPL7428. Used in: Figure 4B |
| pPL7815 | A115L_PMP22-msGFP. drug selection: Amp. | made in pPL7428. Used in: Figure 4B |
| pPL7816 | I166A_PMP22-msGFP. drug selection: Amp. | made in pPL7428. Used in: Figure 4B, 6A, 6B |
| pPL7817 | I137A_PMP22-msGFP. drug selection: Amp. | made in pPL7428. Used in: Figure 4B |
| pPL7818 | V141A_PMP22-msGFP. drug selection: Amp. | made in pPL7428. Used in: Figure 4B |
| pPL7837 | L71A_PMP22-msGFP. drug selection: Amp. | made in pPL7428. Used in: Figure 4B, 6A, 6B |
| pPL7741 | A115V_PMP22-msGFP. drug selection: Amp. | made in pPL7428. Used in: Figure 5B |
| pPL7742 | T118M_PMP22-msGFP. drug selection: Amp. | made in pPL7428. Used in: Figure 5B |
| pPL7743 | I137V_PMP22-msGFP. drug selection: Amp. | made in pPL7428. Used in: Figure 5B |
| pPL7469 | EMP1-msGFP in pcDNA. drug selection: Amp. | pPL7469. Used in: Figure 4A |
| pPL7470 | EMP2-msGFP in pcDNA. drug selection: Amp. | pPL7470. Used in: Figure 4A |
| pPL7628 | JAM_B-HA in pcDNA. drug selection: Amp. | pPL7628. Used in: Figure 6B |
| pPL7629 | JAM_C-HA in pcDNA. drug selection: Amp. | pPL7629. Used in: Figure 6B |
| pPL7896 | EMP1-HA in pcDNA. drug selection: Amp. | pPL7896. Used in: Figure 6A |
| pPL7677 | PMP22-HA in pcDNA3. drug selection: Amp. | pPL7677. Used in: Figure 6A |
| pPL7881 | PMP22_mycEXO in IRES_PacBlue_PB. drug selection: Amp. | pPL7881. Used in: Figure 6C, Suppl.Fig 5, 6 |
| pPL7886 | G150D_PMP22_mycEXO in IRES_PacBlue_PB. drug selection: Amp. | pPL7886. Used in: Figure 6C, Suppl.Fig 5, 6 |
| pPL7885 | L71A_PMP22_mycEXO in IRES_PacBlue_PB. drug selection: Amp. | pPL7886. Used in: Figure 6C, Suppl.Fig 5 |
| pPL7883 | A67T_PMP22_mycEXO in IRES_PacBlue_PB. drug selection: Amp. | pPL7883. Used in: Figure 6C, Suppl.Fig 5, 6 |
| pPL7882 | F75A_PMP22_mycEXO in IRES_PacBlue_PB. drug selection: Amp. | pPL7886. Used in: Figure 6C, Suppl.Fig 5 |
| pPL7884 | I70A_PMP22_mycEXO in IRES_PacBlue_PB. drug selection: Amp. | pPL7886. Used in: Figure 6C, Suppl.Fig 5 |

**Supplemental Figure 1.**
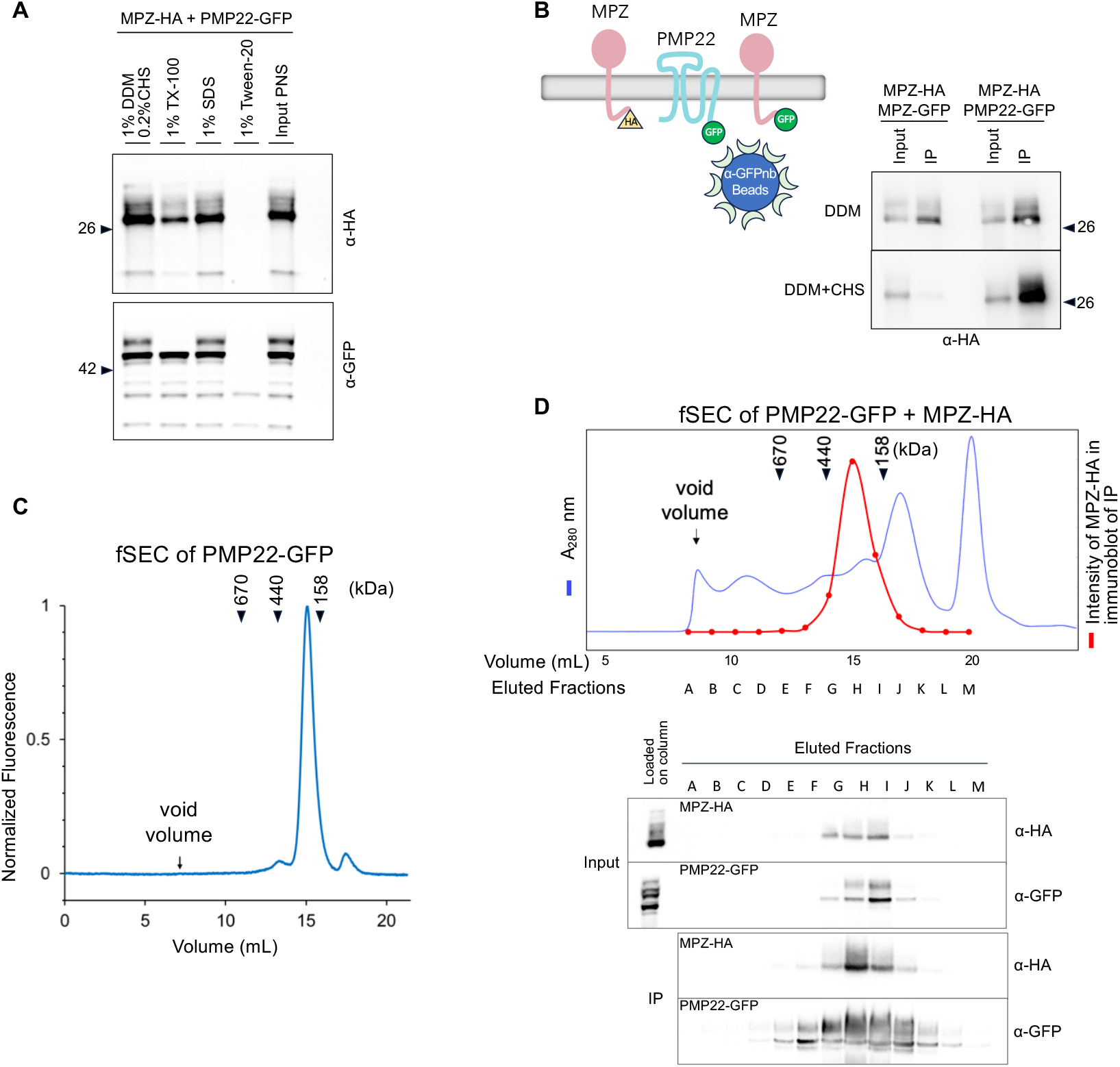
Solubility and Size exclusion chromatography of PMP22-GFP and MPZ-HA. Related to Figure 1. **A**. HEK293 cells expressing MPZ-HA and PMP22-GFP via transient transfection were pelleted, lysed on ice in PBS buffer supplemented with protease inhibitors by passing cells 10 times through syringe with 27-gauge needle. After 20 min, the lysate was centrifuged at 3,000 rpm 5 min to produce a post-nuclear supernatant (PNS). The indicated detergents were added to PNS, incubated 30 min at 20°C and then centrifuged at 200,000xg for 30 min. The volumes of resulting supernatants equivalent to the starting PNS for MPZ-HA and PMP22-GFP were mixed with SOS-sample buffer and heated at 70°C 10 min, then immunoblotted with α-GFP or α-HA. **B**. Co-immunoprecipitations were performed using HEK293 cells expressing MPZ-HA and MPZ-GFP, or MPZ-HA and PMP22-GFP. Proteins in cell lysates were solubilized in either 1% OOM alone or with the addition of 0.2% CHS. GFP fusion proteins were immunoprecipitated with α-GFP nanobody beads, eluted in SOS-sample buffer and immunoblotted with α-GFP or α-HA. **C**. Solubilized lysate from HEK293 cells expressing PMP22-GFP was separated on a Superose 6 10/300 column by HPLC (Shimadzu) and equilibrated with PBS buffer pH 7.4 containing 0.05% OOM and 0.01% CHS, at 0.5 mL/min, and observed by fluorescence detection of the GFP fusion protein (Ex/Em: 488/510 nm). **D**. Solubilized lysate from HEK293 cells expressing PMP22-GFP and MPZ-HA was separated over Superose 6 10/300 column with the same buffer and rate as in C by FPLC (BioRad). Elution fractions (1 mL) after void volume were each immunoprecipitated with α-GFP nanobody beads and then immunoblotted with α-GFP or α-HA. lmmunoblots show distribution of MPZ-HA and PMP22-GFP across the column fractions (noted as lnput) as well as the distribution of the PMP22-GFP~MPZ-HA complex found by MPZ-HA co-immunoprecipitated with PMP22-GFP. Chromatograph shows A280 for total cell lysate protein (blue) and the position of the MPZ-HA~PMP22 complex in elution fractions based on α-HA immunoblot band analysis by lmageJ (red). Approximate sizes (kOa) for elution position of protein markers for the column calibration in PBS buffer shown.

**Supplemental Figure 2.**
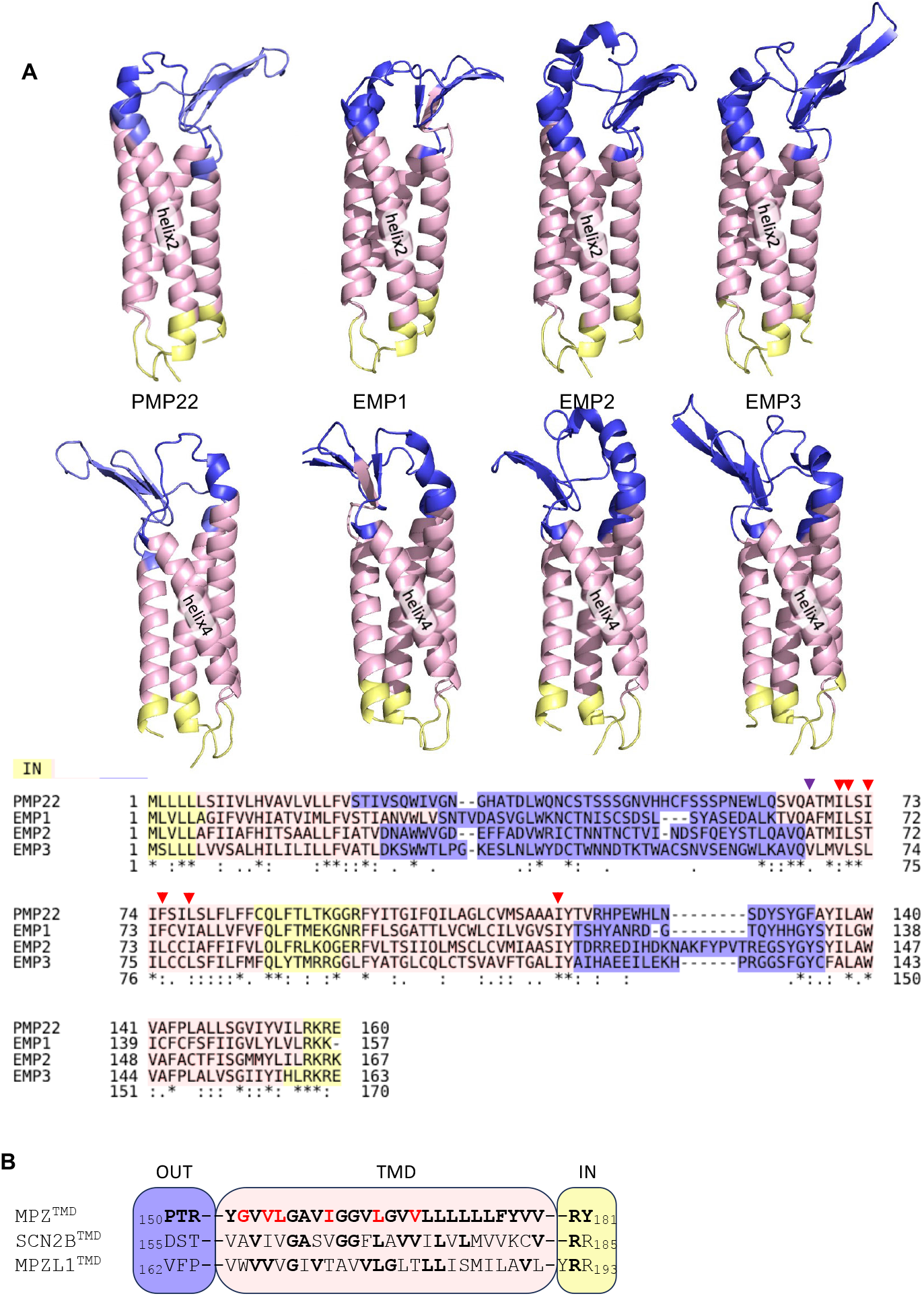
Homology with the MPZ subfamily and PMP22/EMP subfamily of proteins. Related to Figure 4. **A**. Protein sequences for PMP22 and the EMP proteins were aligned using T-Coffee (Tree-based Consistency Objective Function for alignment Evaluation) using the PSI/TM module to predict homology, transmembrane domains, and topology. The various domains (helical transmembrane (pink), cytosolic (yellow), extracellular (blue) are color coded onto structural models from Alphafold2 (Q01453, P54852, P54851, P54849), above. Below, is the sequence alignment. Red arrows denote residues that when altered block MPZ association. The blue arrow points the variant position (A67T) that causes HNPP. **B**. Alignment of the transmembrane domain of MPZ, SCN2B and MPZL1. Residues that when mutated lead to loss of PMP22 association are marked in red. Residues in SCN2B or MPZL1 that are identical in MPZ are shown in bold.

**Supplemental Figure 3.**
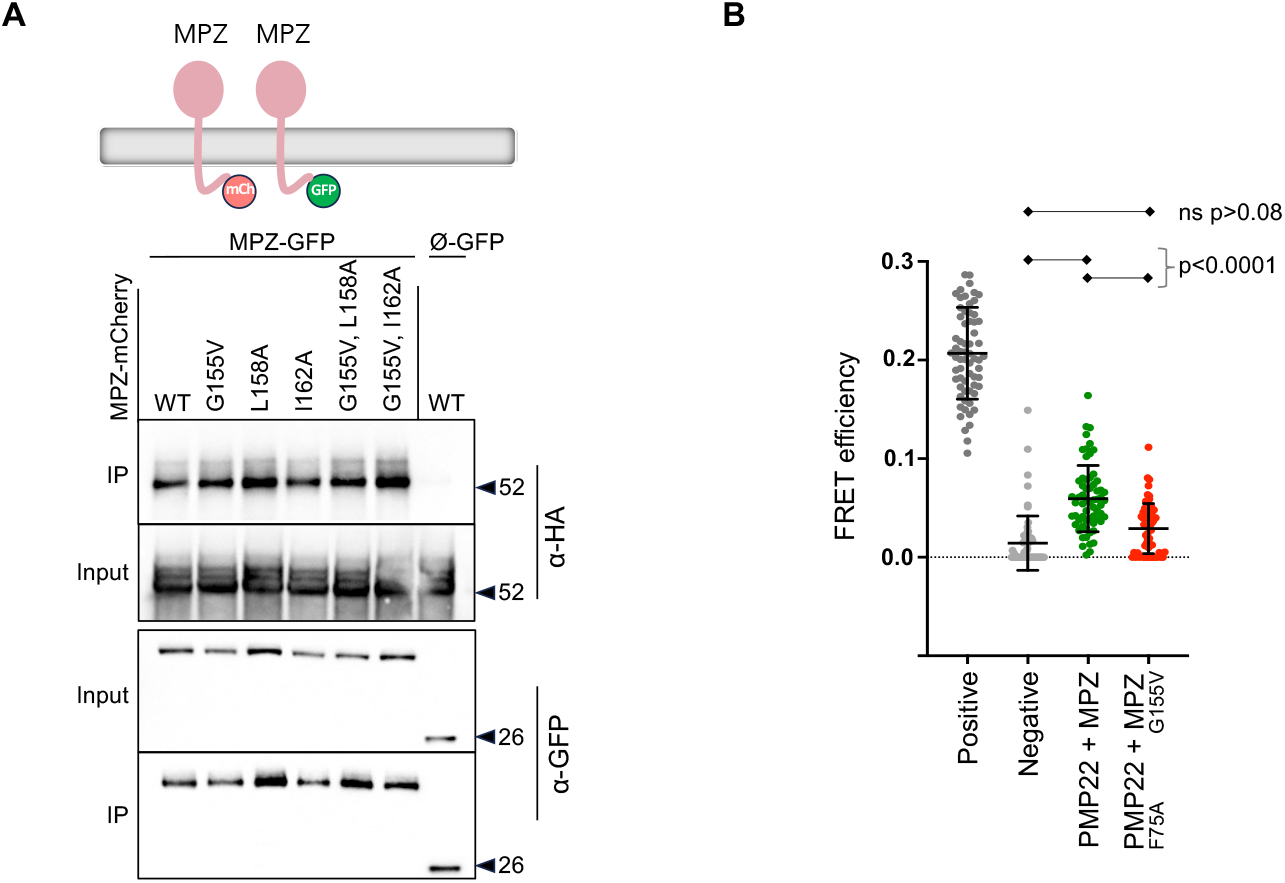
Associations of MPZ-mCherry mutants with MPZ-GFP and PMP22-GFP. Related to Figure 3. **A**. WT and the indicated MPZ-mCherry mutants were assessed for their ability to co-immunoprecipitate with MPZ-GFP using α-GFP nanobody beads. Starting input lysate and immunoprecipitates were immunoblotted with α-mCherry and α-GFP antibodies. **B**. Quantitation of FRET efficiency between PMP22-GFP and MPZ-mCherry and between PMP22-GFP (F75A) and MPZ-mCherry (G155V). FRET was measured as an increase in donor fluorescence after acceptor photobleaching in transiently transfected HEK293 cells. The ‘positive’ control was MPZ tandemly fused to both msGFP2 and mCherry. The ‘negative’ control was Glycophorin A-GFP and MPZ-mCherry. Regions of interest were analyzed and quantified for at least 30 cells from 2 different experiments. FRET efficiencies were calculated with the Leica LAS X Microlab module. All ROI data are represented singly and aggregated as mean FRET efficiency ± S.D. One-way ANOVA with Turkey’s multiple comparison test revealed the negative control was significantly different to each of the other samples.

**Supplementary Figure 4.**
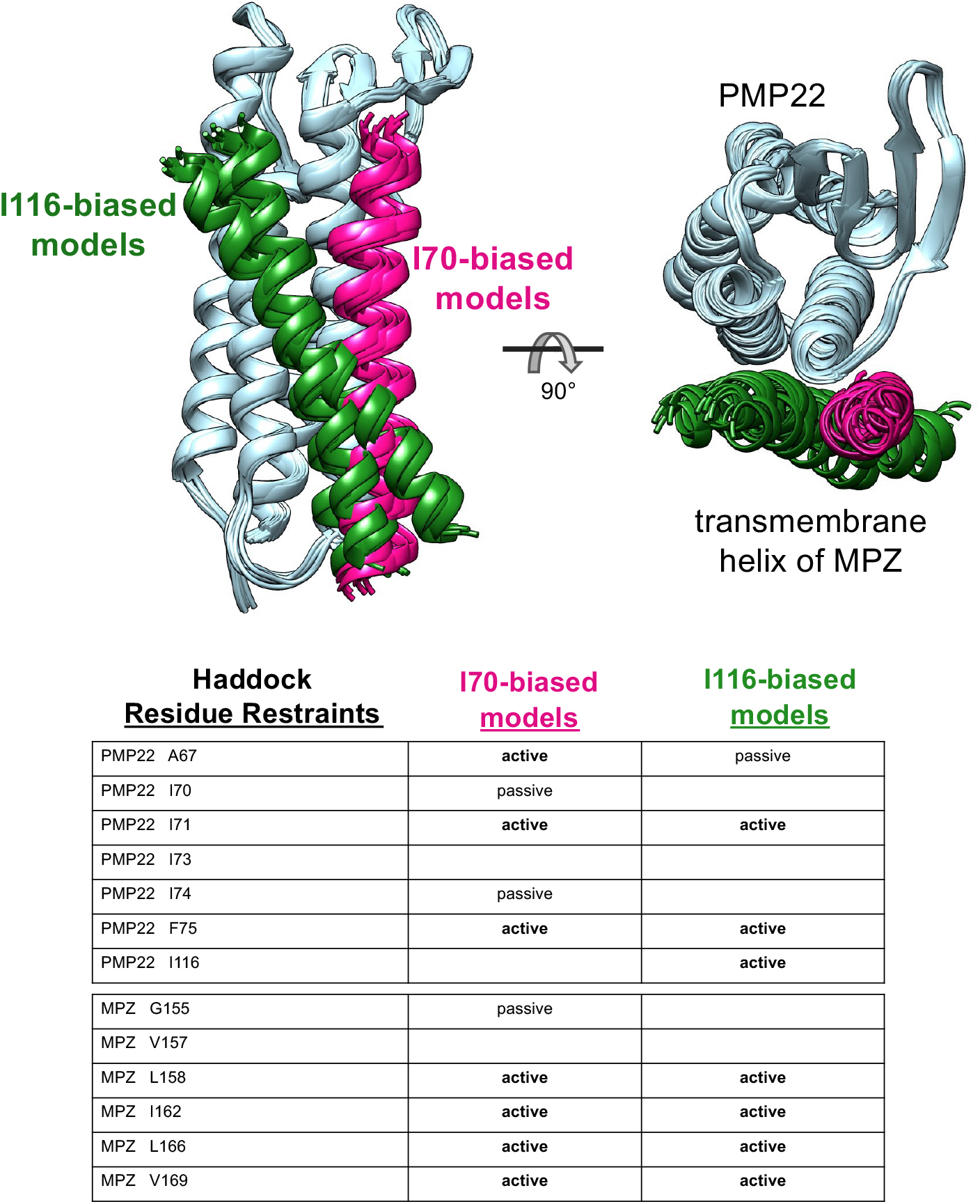
Interaction models of the complex between PMP22 and MPZ. Related to Figure 5. HADDOCK was used to generate a constraint-driven model for the complex between PMP22 (AF-Q01453-F1) and the transmembrane segment of MPZ (AF-P25189-F1). After an initial round of modeling with the full set of HADDOCK residue restraints (listed in table), the resulting feasible poses of model clusters could interact with either I70 or I116 but not both. To limit violations from residues indirectly disrupting the interface, the interaction models were further refined in HADDOCK with the removal of restraints that were irrelevant to the interface for I70- or I116-biased poses. The top 4 models from the one I70-biased model cluster and the two I116-biased model clusters are superimposed on their PMP22 structure (light blue) and depicted with the relative position of the MPZ transmembrane segment (I70-biased – deep pink; I116-biased – forest green). Residue restraints used for the refined HADDOCK docking calculation of biased poses are listed in the table.

**Supplemental Figure 5.**
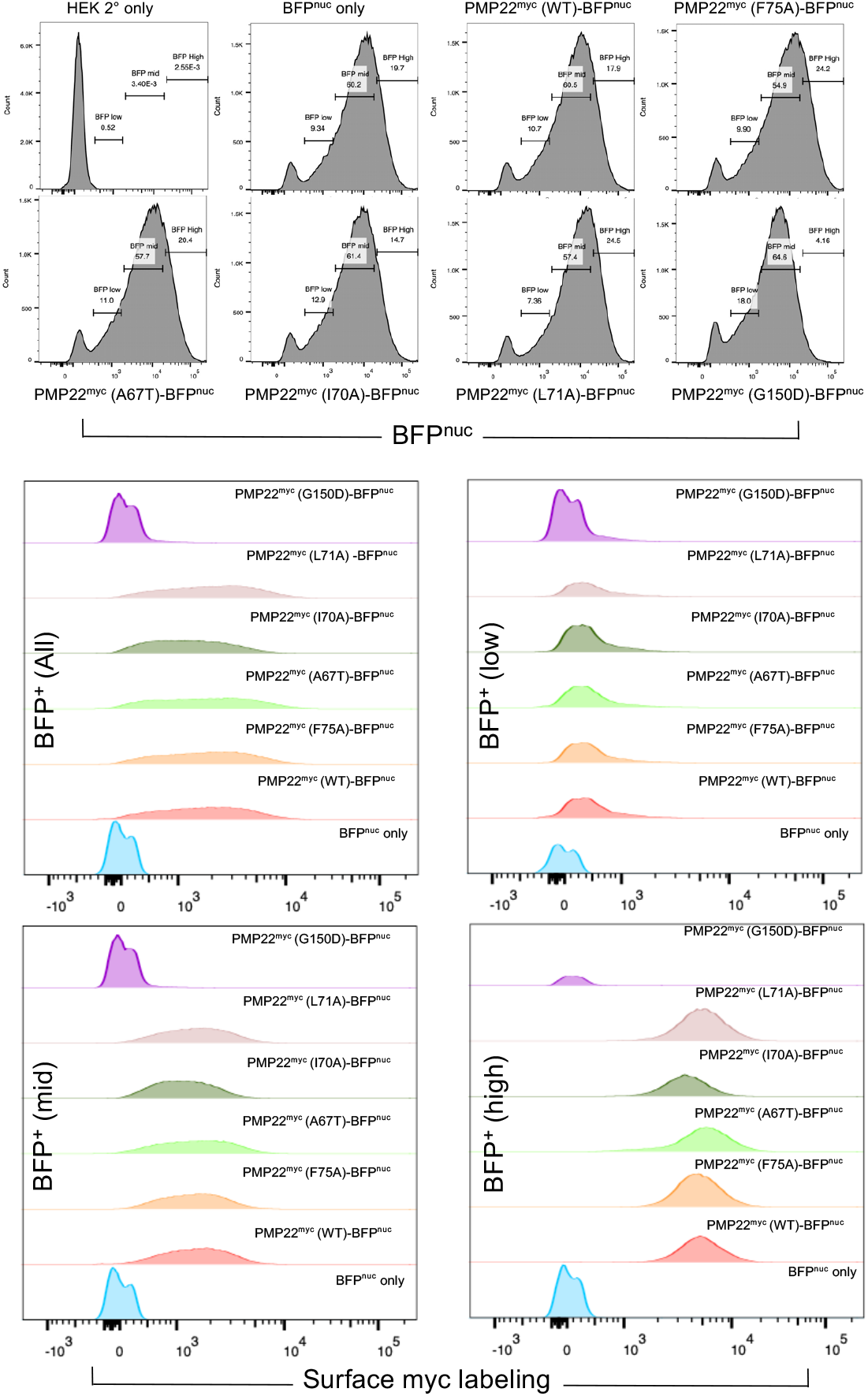
Flow cytometry analysis of PMP22 trafficking to the cell surface. Related to Figure 6. HEK293 cells expressing nuclear localized BFP and the indicated PMP22 mutants carrying an exofacial myc epitope via stable integration of a piggyBac expression plasmid were analyzed by flow cytometry after cell surface labeling with mouse α-myc and goat α-mouse Alexa568 secondary antibody. The top histograms show fluorescence profile of BFP in wildtype cells labeled only with secondary antibody, cells expressing only BFP, or cells expressing BFP and the indicated PMP22-myc^exo^ mutant in tandem. The level of surface myc-tagged protein expression is plotted for the total population as well as 3 subpopulations derived from gating BFP expression from cells expressing low, medium, and high levels of BFP. Since BFP is expressed from the same mRNA as PMP22, the levels of BFP protein are used to normalize expression of PMP22 protein. The bottom images show overlays of PMP22-myc^exo^ on cell surface, presenting comparable surface density of all PMP22 mutants except the PMP22 G150O mutant.

**Supplemental Figure 6.**
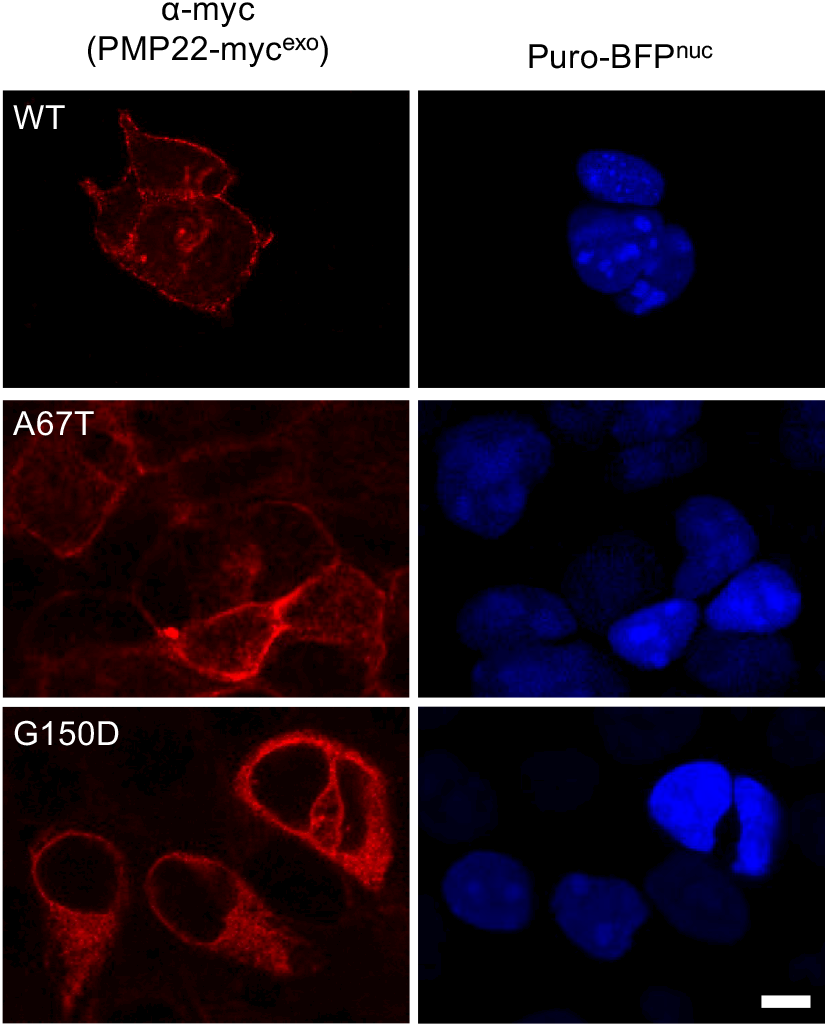
lmmunofluorescence Localization of PMP22myc^exo^ mutants in permeabilized cells. Related to Figure 6. Cell populations described in Figure 6C were fixed, permeabilized in 0.2% Triton X-100, and immunolabelled with α-myc antibodies and Alexa 568-conjugated secondary antibodies (left). Right panels show nuclear localized BFP indicating expression of the indicated PMP22 constructs. Bar −5µm

## Notes

https://figshare.com/articles/dataset/dx_doi_org_10_6084_m9_figshare_6025748/6025748

### Competing Interest Statement

The authors have declared no competing interest.

### Summary of Updates

We provide new data using freeze fracture electron microscopy and quantitative flow cytometry to examine properties of PMP22 and MPZ. We also provide new data showing the chromatographic behavior of the PMP22~MPZ complex. Lastly, we have added clarifying text and a deeper discussion of the data

